# ClairS-TO: A deep-learning method for long-read tumor-only somatic small variant calling

**DOI:** 10.1101/2025.03.10.642523

**Authors:** Lei Chen, Zhenxian Zheng, Junhao Su, Xian Yu, Angel On Ki Wong, Jingcheng Zhang, Yan-Lam Lee, Ruibang Luo

**Author notes:** These authors contributed equally to this work (Lei Chen, Zhenxian Zheng).

## Abstract

Accurate identification of somatic variants in tumor is crucial but challenging, and typically requires a matched normal sample for reliable detection, which is often unavailable in real-world research and clinical scenarios, necessitating proficient algorithms to tell real somatic variants from germline variants and background noises. However, existing tumor-only somatic variant callers that were designed for short-read data don’t work well with long-read. To fill the gap, we present ClairS-TO, a deep-learning-based method for long-read tumor-only somatic variant calling. ClairS-TO uses an ensemble of two disparate neural networks that were trained from the same samples but for opposite tasks – how likely/not likely a candidate is a somatic variant. ClairS-TO also applies multiple post-calling filters, including 1) nine hard-filters, 2) four public plus any number of user-supplied PoNs, and 3) a module that statistically separates somatic and germline variants using tumor purity and copy number profile. Benchmarks using COLO829 and HCC1395 show that ClairS-TO outperforms DeepSomatic in long-read. ClairS-TO is also applicable to short-read and outperforms Mutect2, Octopus, Pisces, and DeepSomatic. Extensive experiments across various sequencing coverages, VAF ranges, and tumor purities support that ClairS-TO has a broad coverage of usage scenarios. ClairS-TO is open-source, available at https://github.com/HKU-BAL/ClairS-TO.

## Introduction

Somatic mutations play a pivotal role in the development and progression of cancer^1^. Accurate identification of the somatic variants is essential for understanding tumorigenesis, developing targeted therapies, and advancing precision oncology^2-4^. Typically, the detection of somatic variants relies on the comparison between tumor and matched normal samples, allowing for identifying somatic mutations from germline polymorphisms^5, 6^. However, in many real-world clinical scenarios, matched normal is not always available^7-11^, necessitating developing tumor-only somatic variant caller.

Tumor-only somatic variant calling is challenging compared with calling in paired samples. Without the matched normal as a reference, distinguishing true somatic variants from germline variants is very challenging. It is difficult to distinguish somatic variants that have a variant allelic fraction (VAF) close to germline variants, let alone the number of germline variants in a sample is often two orders of magnitude more than somatic variants. It is also difficult to distinguish somatic variants with low VAF from background noises (i.e., sequencing errors, alignment artifacts, etc.) without using a paired normal as a reference. It relies heavily on the algorithm itself to tell somatic variants apart from germline variants and background noises when calling somatic variants with tumor sample only. Endeavours have been made to solve the problem for Illumina short-read. Several short-read tumor-only somatic variant callers are commonly used in recent studies^8, 11-14^. These callers primarily use statistical methods that are designed for short-read but inadequate or not applicable to long-read which is known for its relatively higher sequencing error-rate and varying error profile in different challenging genomic regions^15^. Noteworthy, long-read sequencing technologies including Oxford Nanopore Technologies (ONT) and Pacific Biosciences (PacBio) are becoming more relevant for cancer research^5, 6, 16, 17^ and clinical diagnosis^18, 19^ due to their superior performance to span across complex genomic regions and in recognizing disease-causing structural variants (SV). In view of such a trend, efficient and accurate long-read somatic variant callers that use only tumor samples are urged.

In this work, we present ClairS-TO, the first publicly available long-read tumor-only somatic variant caller. ClairS-TO has its name stemmed from ClairS^5^, which requires paired tumor-normal samples for somatic variant calling. To maximize the algorithm’s own ability to tell somatic variant from germline and noises, ClairS-TO leverages an ensemble of two disparate neural networks that were trained from the same samples but for two exact opposite tasks – an affirmative neural network that determines how likely a candidate is a somatic variant, and a negational neural network that determines how likely a candidate is not a somatic variant. A posterior probability for each variant candidate is calculated from the outputs of the two networks and prior probabilities derived from the training samples. After the networks, ClairS-TO applies three techniques to further remove non-somatic variants: 1) Nine hard-filters that were found effective for short-read are applied but with algorithms optimized or parameters tuned for long-read; 2) Four panels of normals (PoNs), including three built from short-read datasets and one from long-read datasets are applied; and 3) a statistical method is applied to classify each variant into either a germline, somatic, or a subclonal somatic variant, using the estimated tumor sample purity and ploidy, as well as the copy number profile of each variant. About model training strategy, ClairS-TO depends on synthetic tumor samples, and can be augmented using real tumor samples. Synthetic tumor sample combines the real sequencing reads of two samples and regards a germline variant that is specific to one sample as a somatic variant to the other sample. While somatic variants are scarce in real samples, the method can generate a number of training samples comparable to that of germline variants, providing sufficient samples to train a deep neural network robustly. The method also enabled fine-tuning the network using a much smaller number of somatic variants available in the real tumor samples, which is essential to allow the network to learn cancer-specific variant characteristics, such as mutational signatures^20^.

We evaluated the performance of ClairS-TO using COLO829 (metastatic melanoma)^21^ and HCC1395 (breast cancer)^22^. Both cancer samples are richly studied and with reliable truths for benchmarking. With ONT Q20+ long-reads, our experiments show that ClairS-TO consistently outperformed DeepSomatic (another long-read tumor-only somatic variant caller released after ClairS-TO) at multiple coverages, tumor purities, and VAF ranges. With PacBio Revio long-reads, ClairS-TO also outperformed DeepSomatic but with a smaller edge. ClairS-TO is optimized for long-read but is also applicable to short-read. Our experiments also show that ClairS-TO has outperformed Mutect2, Octopus, Pisces, and DeepSomatic at 50-fold coverage of Illumina short-reads.

## Results

### Overview of ClairS-TO

ClairS-TO comprises two disparate neural networks, the affirmative network (AFF) and the negational network (NEG), and is specifically designed for long-read tumor-only somatic variant calling. ClairS-TO offers two pre-trained models for inference: 1) a model trained exclusively on **s**ynthetic **s**amples (ClairS-TO SS), and 2) a model initially trained on **s**ynthetic **s**amples and further augmented with **r**eal **s**amples (ClairS-TO SSRS), as illustrated in **Figure 2a**. For the SS model training, synthetic tumors were typically created by mixing two biologically unrelated GIAB samples^23^. ClairS-TO SSRS was then fine-tuned using the SS model weights with real cancer cell lines, as detailed in the **Method** – **Training data preparation** section. **Figure 1a** provides an overview of the ClairS-TO somatic variant calling workflow. Following the ensembled network prediction, ClairS-TO applies three post-filtering steps: artifact filtering with nine hard-filters, germline variant tagging with four PoNs, and the Verdict module for distinguishing germline and somatic variants. Further details are described in the **Method** – **ClairS-TO workflow and design** section.

**Figure 1.**
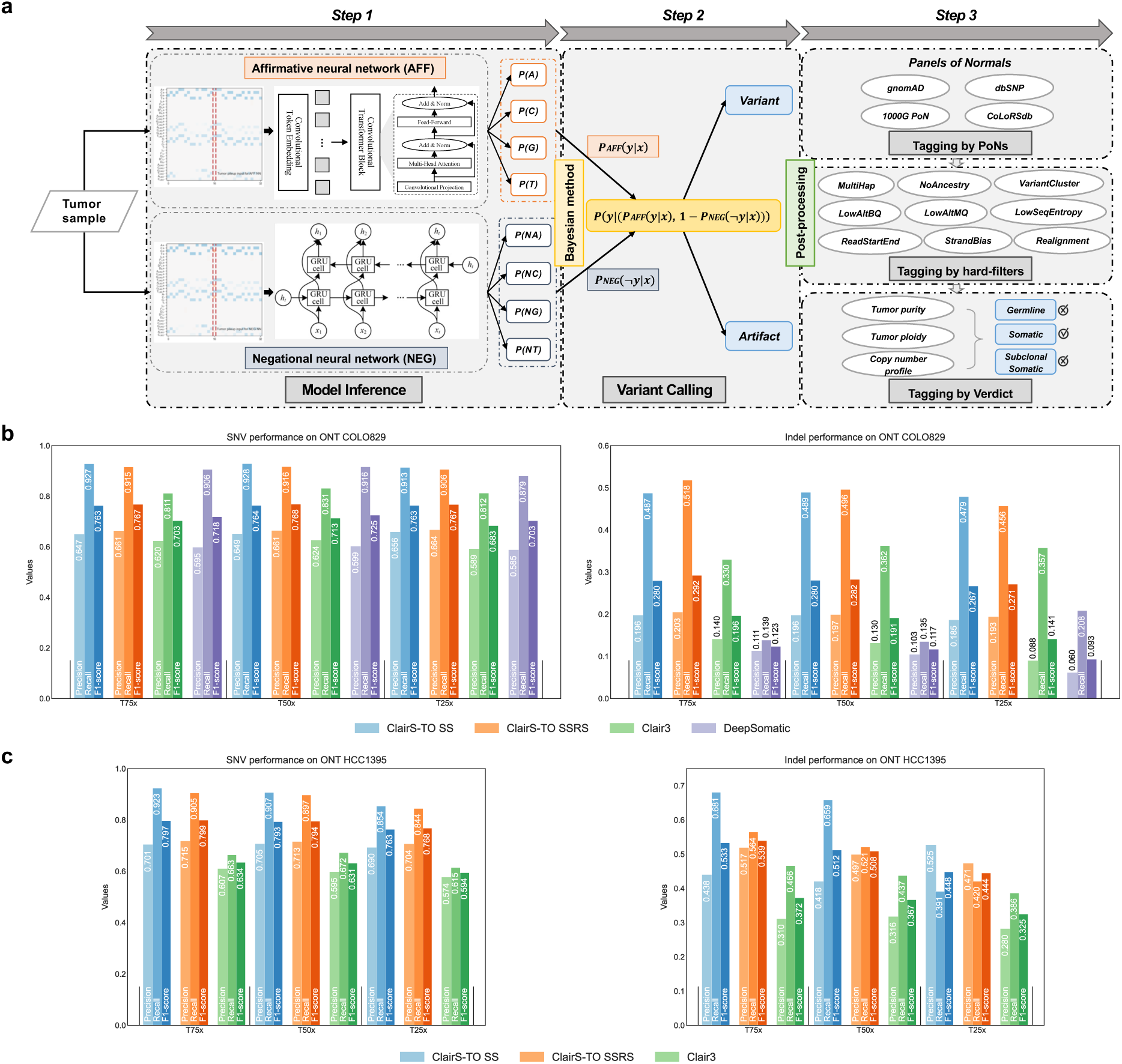
Overview of ClairS-TO somatic variant calling workflow. (a) The three steps in the somatic variant calling workflow of ClairS-TO. In Step 1, the pileup inputs of variant candidates in the tumor are fed into the affirmative neural network (AFF) and the negational neural network (NEG) to obtain the probabilities of the candidate being a somatic variant and not being a somatic variant, respectively. In Step 2, a joint posterior probability *P*(*y*|(*P*_*AFF*_(*y*|*x*), 1 − *P*_*NEG*_(¬*y*|*x*))) is calculated using the output probabilities from both networks for inference. *P*(*y*) is derived from the same samples used for training AFF and NEG. In Step 3, multiple post-processing strategies are applied to tag non-somatic variants, including: 1) leveraging PoNs to tag germline variants, 2) applying nine hard-filters to remove sequencing artifacts, and 3) using the Verdict module to further distinguish somatic variants from germline variants by modeling tumor sample purity, tumor sample ploidy, and the copy number profile. (b) The performance of the best achievable F1-score, along with the corresponding precision and recall, at 25-, 50-, and 75-fold tumor coverages of ONT COLO829. (c) The performance of the best achievable F1-score, along with the corresponding precision and recall, at 25-, 50-, and 75-fold tumor coverages of ONT HCC1395. DeepSomatic was excluded from benchmarking with the HCC1395 WGS dataset because the sample was included in its model training.

**Figure 2.**
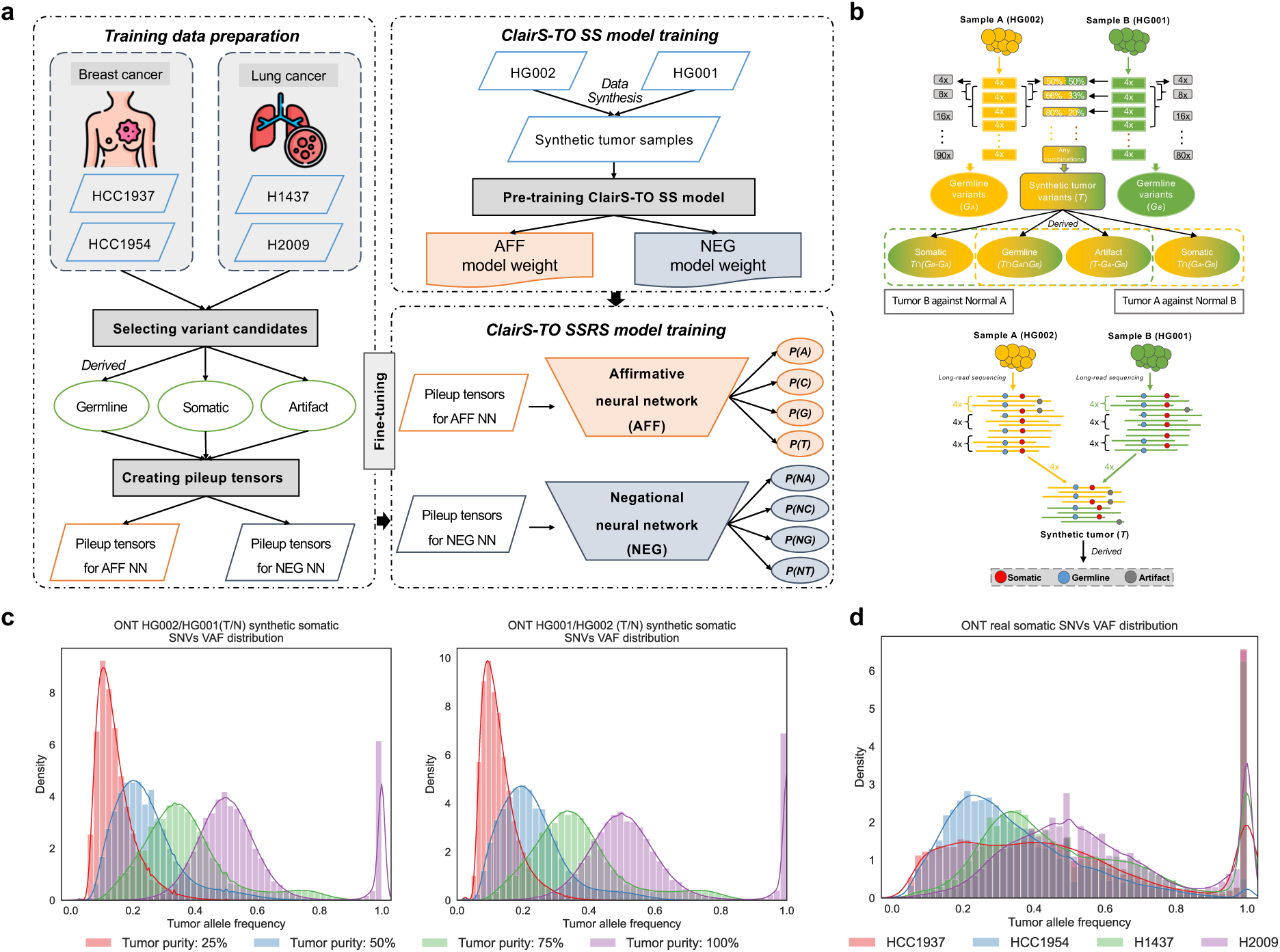
Overview of ClairS-TO training workflow. (a) The training workflow of ClairS-TO. ClairS-TO provides two types of pre-trained models: a model trained exclusively on synthetic tumor samples (ClairS-TO SS) and a model initially trained on synthetic tumor samples and then augmented with real tumor samples (ClairS-TO SSRS). For ONT, the ClairS-TO SS model was trained using synthetic tumors generated from GIAB HG001 and HG002. ClairS-TO SSRS was fine-tuned from the ClairS-TO SS model using four real cancer cell lines (HCC1937, HCC1954, H1437, and H2009). (b) The data synthesis workflow of ClairS-TO. For example, using 90-fold coverage of GIAB HG002 (Sample A, with GIAB-known truth germline variants as G_A_) and 80-fold coverage of GIAB HG001 (Sample B, with GIAB-known truth germline variants as G_B_) from ONT WGS alignments as sources for synthetic tumor and normal, the alignments were split into smaller non-overlapping chunks with an average of 4-fold coverage. These smaller chunks were used to simulate synthetic tumors with varying coverages and different VAFs. For variants in the synthetic tumor (T), "Somatic" refers to GIAB truth germline variants present in A but not in B (T∩(G_A_-G_B_)); "Germline" refers to GIAB truth germline variants present in both A and B (T∩G_A_∩G_B_); and "Artifact" refers to variants not present in the GIAB truth germline variants of either A or B (T-G_A_-G_B_). When using Sample B as the tumor and Sample A as the normal, the definitions remain identical except for switching the subscripts. (c) The VAF distributions of synthetic somatic SNVs at simulated tumor purities of 100%, 75%, 50%, and 25%, using either HG002 or HG001 as the tumor source. (d) The VAF distributions of real somatic SNVs in the four real cancer cell lines (i.e., HCC1937, HCC1954, H1437, and H2009).

### Performance analysis on ONT

A summary of the ONT datasets used for model training and benchmarking is provided in **Supplementary Table 1a**. ClairS-TO SS was trained on synthetic datasets derived from GIAB HG002 and HG001 from the ONT EPI2ME Labs^24^. ClairS-TO SSRS was augmented from the SS model using four real cancer cell lines – HCC1937, HCC1954, H1437, and H2009 – from Park et al.^6^. For performance evaluation, two well-established cancer cell lines, COLO829 and HCC1395, were used. Our evaluation involved comparing results with other callers, including Clair3^25^ and DeepSomatic^6^. Since Clair3 is a germline variant caller, we treated variants called by Clair3 as somatic, excluding those tagged by the PoNs used in ClairS-TO. DeepSomatic is a deep-learning-based somatic variant caller that supports tumor-only calling and is trained exclusively on real cancer cell lines. For benchmarking, we used the best-performing "multi-cancer" model, following the authors’ recommended practices.

#### Performance with different sequencing coverages

We evaluated ClairS-TO and other callers across three sequencing coverages: 25-, 50-, and 75-fold, considering that 25-fold coverage typically represents the throughput of one ONT R10.4.1 PromethION flow cell. We incrementally increased the coverage by one flow cell throughput, aligning with clinical practices to enhance variant discovery capacity.

**Figure 3a** and **Supplementary Table 2** showed the AUPRC (area under precision-recall curve) and the best achievable F1-score results of each caller. For the COLO829 dataset, ClairS-TO SSRS achieved 0.6489, 0.6634, and 0.6685 AUPRC for SNV at 25-, 50-, and 75-fold coverage, respectively. These values were 0.0336, 0.0458, and 0.0494 higher than those of the SS model. The improvement is more pronounced from 25- to 50-fold (+0.0145 AUPRC) compared with 50- to 75-fold (+0.0051 AUPRC). The SSRS model’s consistent outperformance against the SS model demonstrates that incorporating real samples during training enhances the ability to learn variant characteristics inherent in real tumors.

**Figure 3.**
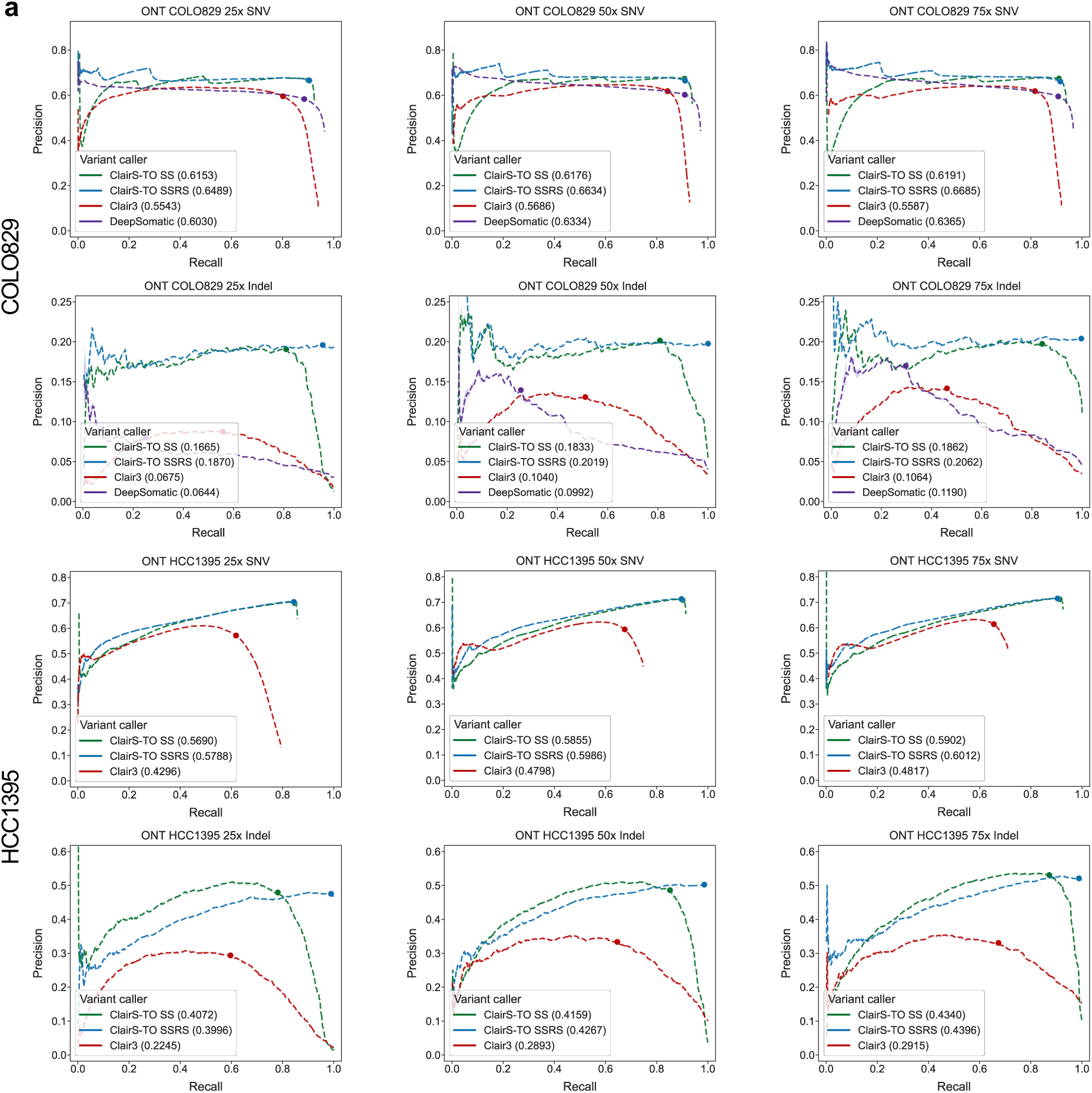
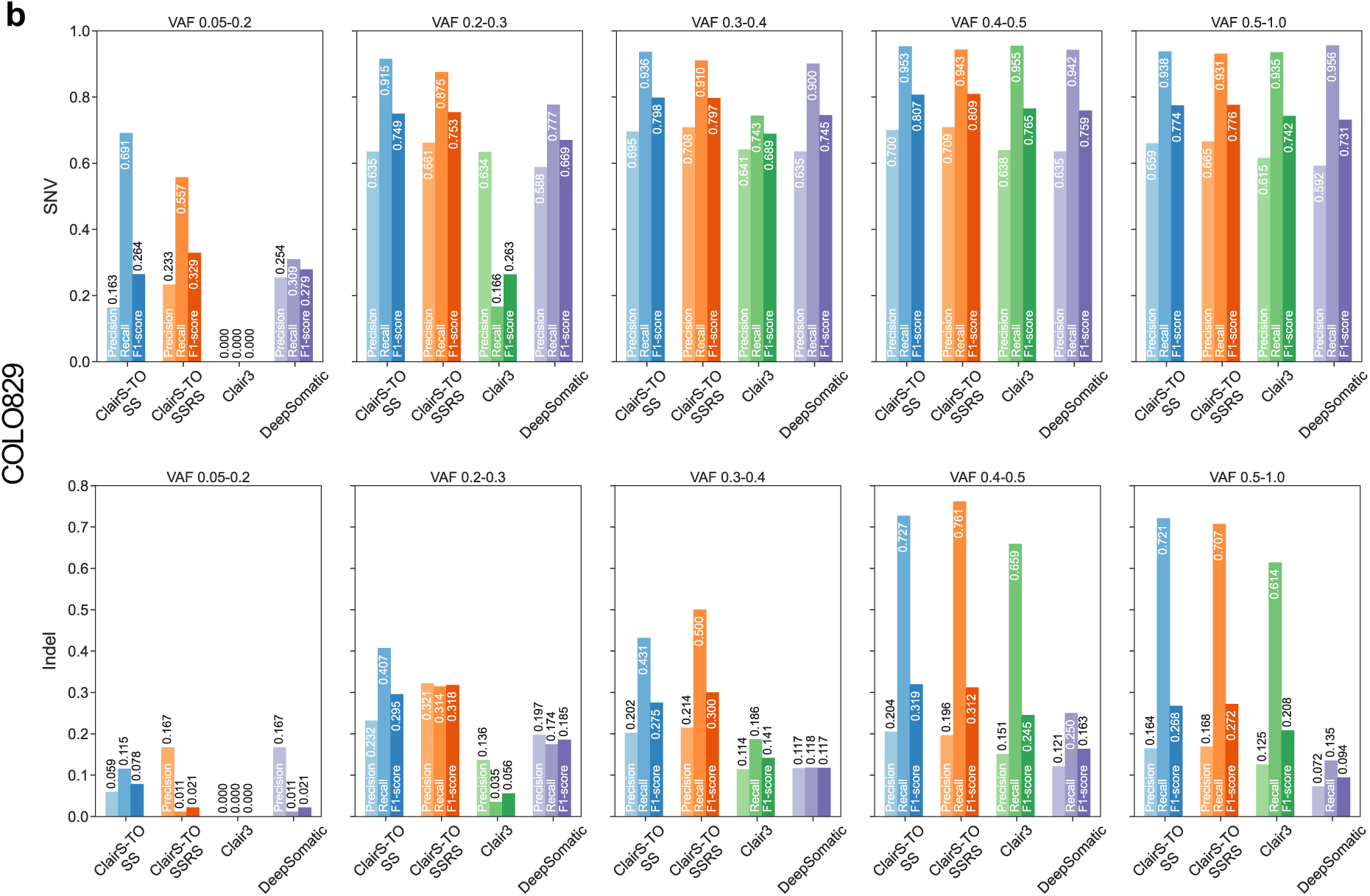
Performance evaluation at different sequencing coverages and VAF ranges. a) Precision-Recall curves at 25-, 50-, and 75-fold tumor coverages of ONT COLO829 and HCC1395. The dot on each dashed line indicates where the best F1-score was achieved. The area under the Precision-Recall curve (AUPRC) is shown in parentheses in the legend. (b) Performance on 50-fold coverage of ONT COLO829 across different VAF ranges, with VAF divided into five ranges: 0.05–0.2, 0.2–0.3, 0.3–0.4, 0.4–0.5, and 0.5–1.0.

Among other callers, Clair3 achieved AUPRC values of 0.5543, 0.5686, and 0.5587 at the three coverages, underscoring the limitations of using a germline variant caller for somatic variant calling. DeepSomatic achieved AUPRC values of 0.6030, 0.6334, and 0.6365 at the three coverages, which were 0.0459, 0.0300, and 0.0320 lower than those of ClairS-TO SSRS, respectively. In terms of the best achievable F1-score, as shown in **Figure 1b**, ClairS-TO SSRS achieved 76.66%, 76.83%, and 76.75% at the three coverages, surpassing DeepSomatic by 6.38%, 4.38%, and 4.91%, respectively. This improvement highlights the advantage of augmenting training with both synthetic and real tumor samples.

For the HCC1395 dataset, DeepSomatic was excluded from benchmarking because HCC1395 was part of its training data. The rationale for ClairS-TO’s decision to exclude HCC1395 entirely, and how this decision benefits the community by enabling more comprehensive performance comparisons between algorithms and tools in the long term, is explained in the **Method** – **Training data preparation** section. As shown in **Figure 3a** and **Figure 1c**, ClairS-TO SSRS performed best, achieving an average AUPRC of 0.5929 and an F1-score of 78.69% across the three coverages. The results across different datasets demonstrate that ClairS-TO consistently outperforms other callers under various coverage settings. Notably, ClairS-TO achieved ∼92% recall across different coverages, comparable to ClairS, a tumor-normal-pair caller, making ClairS-TO a reliable option for identifying somatic variants of interest with clinical sequencing panels.

#### Performance at different VAF ranges

We also benchmarked ClairS-TO across different VAF ranges using 50-fold coverage of COLO829, as shown in **Figure 3b** and **Supplementary Table 3**. In the low VAF range (0.05–0.2), low-mid VAF ranges (0.2–0.3, 0.3–0.4, and 0.4–0.5), and mid-high VAF range (0.5–1.0), ClairS-TO SS achieved F1-scores of 26.38%, 74.94%, 79.79%, 80.67%, and 77.43%, respectively. In comparison, ClairS-TO SSRS achieved F1-scores of 32.85%, 75.35%, 79.66%, 80.90%, and 77.60%. Across all VAF ranges, ClairS-TO consistently outperformed Clair3 and DeepSomatic, as detailed in **Supplementary Table 3**. Notably, the performance at the low-mid VAF ranges is similar to the mid-high VAF ranges. While higher VAF ranges typically yield better results, the inclusion of incorrectly called germline variants in the mid-high VAF ranges led to a drop in precision.

#### Performance at different tumor purities

In real-world clinical scenarios, tumor samples are often impure and sequenced with varying tumor purities. We evaluated the performance of ClairS-TO at different tumor purities – 1.0, 0.8, 0.6, 0.4, and 0.2 – by mixing normal samples into tumors *in silico*. The results for 50-fold coverage of COLO829 are shown in **Figure 4a** and **Supplementary Table 4a**. ClairS-TO SSRS achieved AUPRC values of 0.6634, 0.6256, 0.6041, 0.5500, and 0.4797 at the five tumor purities, respectively, while ClairS-TO SS achieved AUPRC values of 0.6176, 0.5930, 0.5849, 0.5470, and 0.4851. ClairS-TO SSRS outperformed DeepSomatic by 0.0300 to 0.1919 across the range of tumor purities. As shown in **Figure 4b**, ClairS-TO SS/SSRS achieved F1-scores of 76.36%/76.83%, 75.13%/75.86%, 74.88%/75.14%, 74.79%/74.95%, and 66.87%/66.99% at the five tumor purities, respectively. The performance decline was most pronounced when tumor purity dropped from 0.4 to 0.2, primarily due to a reduction in recall.

**Figure 4.**
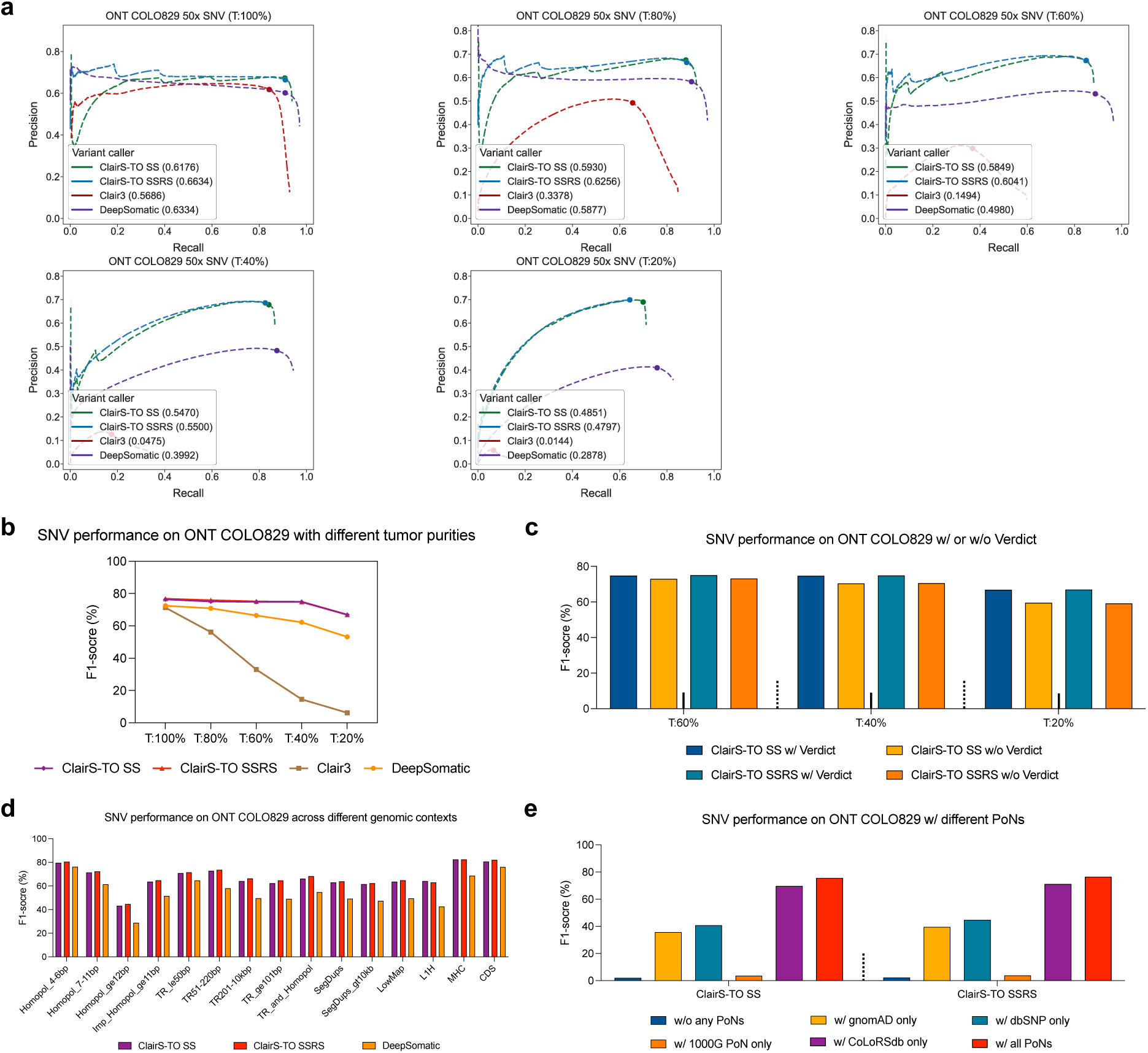
Performance evaluation at different tumor purities, genomic contexts, and program settings. (a) Precision-Recall curves on 50-fold coverage of ONT COLO829 at five tumor purities: 1.0, 0.8, 0.6, 0.4, and 0.2. The dot on each dashed line indicates where the best F1-score was achieved. AUPRC is shown in parentheses in the legend. (b) The best achievable F1-score on 50-fold coverage of ONT COLO829 at five tumor purities: 1.0, 0.8, 0.6, 0.4, and 0.2. (c) Performance with or without the Verdict module. Notably, Verdict significantly improved performance, particularly when tumor purity is low, and is recommended to be enabled when the estimated purity is below 0.6. (d) The best achievable F1-score on 50-fold coverage of ONT COLO829 across different challenging genomic contexts, categorized into five groups: "Low complexity", "Segmental duplications", "Low mappability", "Functional regions", and "Other difficult regions", as defined by GIAB Stratifications v3.3. (e) The best achievable F1-score on 50-fold coverage of ONT COLO829 using different PoNs or combinations for filtering.

A statistical method we named Verdict is used to classify variants as somatic or non-somatic using estimated tumor purity, tumor ploidy, and copy number profile, with further details provided in the **Method** – **ClairS-TO workflow and design** section. As shown in **Figure 4c** and **Supplementary Table 4b**, ClairS-TO SS/SSRS achieved F1-scores of 74.88%/75.14%, 74.79%/74.95%, and 66.87%/66.99% at tumor purities of 0.6, 0.4, and 0.2, respectively, when using the Verdict module. Disabling Verdict resulted in F1-score decreases by 1.89%/1.99%, 4.36%/4.38%, and 7.31%/7.81%. Notably, when tumor purity exceeds 0.6, the improvements are negligible. The results indicate that the Verdict module significantly reduces the misclassification of germline variants, particularly in tumor samples with low purity, even without a matched normal sample as a reference.

#### Performance across various genomic contexts

We benchmarked ClairS-TO using 50-fold coverage of COLO829 and categorized challenging genomic contexts into five categories: "Low complexity", "Segmental duplications", "Low mappability", "Functional regions", and "Other difficult regions", as defined by GIAB Stratifications v3.3^23^. The best achievable F1-scores are shown in **Figure 4d** and **Supplementary Table 5**. ClairS-TO consistently outperformed DeepSomatic across all genomic contexts. Specifically, ClairS-TO SS, ClairS-TO SSRS, and DeepSomatic achieved F1-scores of 80.65%, 82.10%, and 76.16% in coding sequence (CDS) regions, respectively.

Although ClairS-TO outperformed DeepSomatic by 11.11% and 13.69% in complex homopolymer and tandem repeat regions, the best F1-scores achieved by ClairS-TO SSRS were only 65.63% and 69.09%, respectively. These scores remain significantly lower than the whole-genome sequencing (WGS) F1-score (67.36% vs. 76.83%), underscoring the ongoing challenges in somatic variant detection within complex genomic contexts.

#### Benchmarking different network and training configurations

We evaluated the performance of various network configurations on 50-fold coverage of COLO829 using: 1) the default ensemble (AFF & NEG) in ClairS-TO, 2) the AFF network only (AFF only), and 3) the NEG network only (NEG only). As shown in **Supplementary Figure 1b and Supplementary Table 7b**, the ClairS-TO SSRS model achieved an AUPRC of 0.6634 with the AFF & NEG ensemble, while AFF only achieved 0.6363 AUPRC, and NEG only achieved 0.6562 AUPRC. These results highlight the benefits of network ensembling for enhanced performance.

We also compared the performance of three ClairS-TO models: ClairS-TO SS, ClairS-TO RS, and ClairS-TO SSRS, using 50-fold coverage of COLO829. Specifically, ClairS-TO RS was trained from scratch using four real cancer cell lines (HCC1937, HCC1954, H1437, and H2009). As shown in **Supplementary Figure 1a** and **Supplementary Table 7a**, ClairS-TO SS, ClairS-TO RS, and ClairS-TO SSRS achieved AUPRC values of 0.6176, 0.6582, and 0.6634, respectively. The superior performance of ClairS-TO SSRS demonstrates the importance of combining synthetic samples and real tumors for augmented training.

#### Analysis of introducing different PoNs for germline tagging

ClairS-TO utilizes four PoNs – gnomAD^26^, dbSNP^27^, 1000G PoN^28^, and CoLoRSdb^29^ – to tag non-somatic variants. As shown in **Figure 4e** and **Supplementary Table 7c**, using 50-fold coverage of COLO829, ClairS-TO SSRS achieved F1-scores of 2.31%, 39.56%, 44.80%, 3.87%, 71.19%, and 76.49% when using no PoNs, gnomAD only, dbSNP only, 1000G PoN only, CoLoRSdb only, and all four PoNs, respectively. Precision improved by 62.86% when transitioning from no PoNs to using all four PoNs, with a total of ∼3.37 million germline variants tagged. These results underscore the benefits of incorporating multiple PoNs, particularly CoLoRSdb, a PoN specifically designed for long-read data^29^.

#### Performance evaluation of different hard-filters

We further assessed the impact of nine hard-filters, the details of which are described in the **Method** – **ClairS-TO workflow and design** section. Using 50-fold coverage of COLO829, the filters "MultiHap", "NoAncestry", "LowAltBQ", "LowAltMQ", "VariantCluster", "ReadStartEnd", and "StrandBias" removed 125, 26, 30, 1, 6,541, 87, and 237 non-somatic variants, respectively. With these hard-filters applied, precision improved by 1.85%, demonstrating their effectiveness in eliminating germline variants and sequencing artifacts.

### Performance analysis on PacBio

To validate the effectiveness of our training workflow and proposed post-filters, we also trained models on PacBio data, with a summary of the datasets used provided in **Supplementary Table 1a**. The ClairS-TO SS PacBio model was trained on a synthetic dataset derived from GIAB HG003 and HG004, sequenced on the PacBio Revio system. The ClairS-TO SSRS model was augmented from the SS model using four real cancer cell lines (HCC1937, HCC1954, H1437, and H2009) provided by Park et al.^6^. For benchmarking, we used 50-fold coverage of the PacBio COLO829 dataset. As shown in **Figure 5a** and **Supplementary Table 8a**, ClairS-TO SS achieved an AUPRC of 0.6439 and an F1-score of 78.50%, while ClairS-TO SSRS achieved an AUPRC of 0.6667 and an F1-score of 78.64%. In comparison, DeepSomatic achieved an AUPRC of 0.6588 and an F1-score of 74.65%, which is ∼4% lower than ClairS-TO SSRS in F1-score.

**Figure 5.**
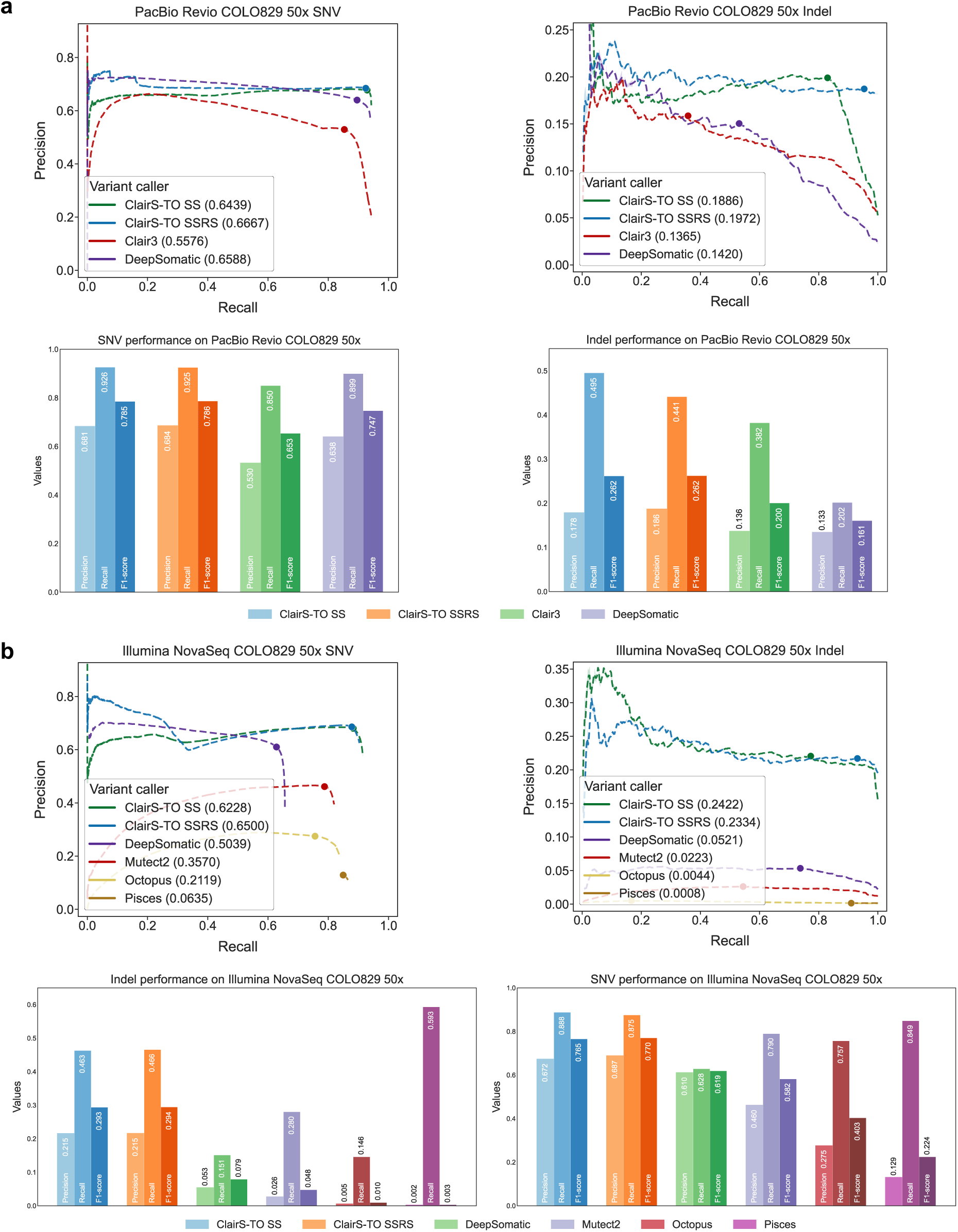
Performance analysis on PacBio and Illumina data. (a) The Precision-Recall curves and the best achievable F1-score of SNV and Indel on coverage PacBio Revio COLO829 using ClairS-TO SS, ClairS-TO SSRS, Clair3, and DeepSomatic. The dot on each dashed line represents the best F1-score achieved. AUPRC is shown in parentheses in the legend. (b) The Precision-Recall curves and the best achievable F1-score of SNV and Indel on 50-fold coverage of Illumina COLO829 using ClairS-TO SS, ClairS-TO SSRS, DeepSomatic, Mutect2, Octopus, and Pisces. The dot on each dashed line represents the best F1-score achieved. AUPRC is shown in parentheses in the legend.

### Performance analysis on Illumina

Short-read tumor-only somatic variant calling has been extensively studied^7-14, 30-32^, with numerous notable methods developed, such as Mutect2^12^, Octopus^13^, and Pisces^14^. ClairS-TO was primarily designed for long-read data but can be adapted to short-read data by excluding long-read-specific post-filters, such as long-read PoNs, variant phasing, and the "MultiHap" and "NoAncestry" filters. For training, we used GIAB HG003 and HG004 for the SS model and augmented the SSRS model with four real cancer cell lines (HCC1937, HCC1954, H1437, and H2009). All datasets were sequenced on Illumina’s latest HiSeq X or NovaSeq 6000 platforms, with more details provided in **Supplementary Table 1a**.

**Figure 5b** and **Supplementary Table 8b** showed the results of ClairS-TO compared to other short-read tumor-only somatic variant callers. ClairS-TO SS/SSRS achieved AUPRC values of 0.6228/0.6500 and F1-scores of 76.50%/76.99% at 50-fold coverage of COLO829, respectively. DeepSomatic ranked second, achieving an AUPRC of 0.5039 and an F1-score of 61.89%, which is 15.10% lower than ClairS-TO SSRS in F1-score. ClairS-TO also outperformed other statistical short-read callers, including Mutect2, Octopus, and Pisces, with F1-score improvements of 18.83%, 36.69%, and 54.59%, respectively. These results demonstrate that ClairS-TO achieves competitive performance on short-read data and could serve as a supplementary tool for ensemble analysis in short-read workflows.

### Indel performance analysis

Indel variant detection is more challenging compared to SNVs, primarily due to two reasons: 1) Long-read technologies exhibit higher error-rate in complex regions, such as homopolymer and tandem repeat regions, leading to substantial Indel artifacts that hinder the detection of true somatic Indels with low VAF; and 2) The number of somatic Indels is inherently limited. For instance, only around 2,000 somatic Indels are reported in HCC1395 or COLO829, resulting in an insufficient number of Indels from real samples for comprehensive model training and benchmarking currently. While tumor-only somatic Indel detection performance remains limited, we provide the following results as a baseline for future improvements.

As shown in **Figure 3a**, **Figure 5a**, and **Figure 5b**, ClairS-TO SS/SSRS achieved AUPRC values of 0.1833/0.2019, 0.1886/0.1972, and 0.2422/0.2334 on ONT, PacBio, and Illumina 50-fold coverage of the COLO829 dataset, respectively. The best achievable F1-scores were 28.24%, 26.21%, and 29.43% across the three platforms. ClairS-TO consistently outperformed DeepSomatic and other short-read callers, as detailed in **Supplementary Table 2**, with similar trends observed in benchmarking at different coverages or using the HCC1395 cell line. Despite ClairS-TO performing the best among the evaluated tools, recall remains capped at ∼50%, highlighting the need for further improvements to achieve reliable somatic Indel detection.

### Analysis of false positive and false negative calls

We manually analyzed 300 false positive (FP) calls and 300 false negative (FN) calls randomly selected from 50-fold coverage of ONT COLO829. Each FP and FN was assigned with a primary and a secondary reason, as shown in **Figure 6** and **Supplementary Table 6**.

**Figure 6.**
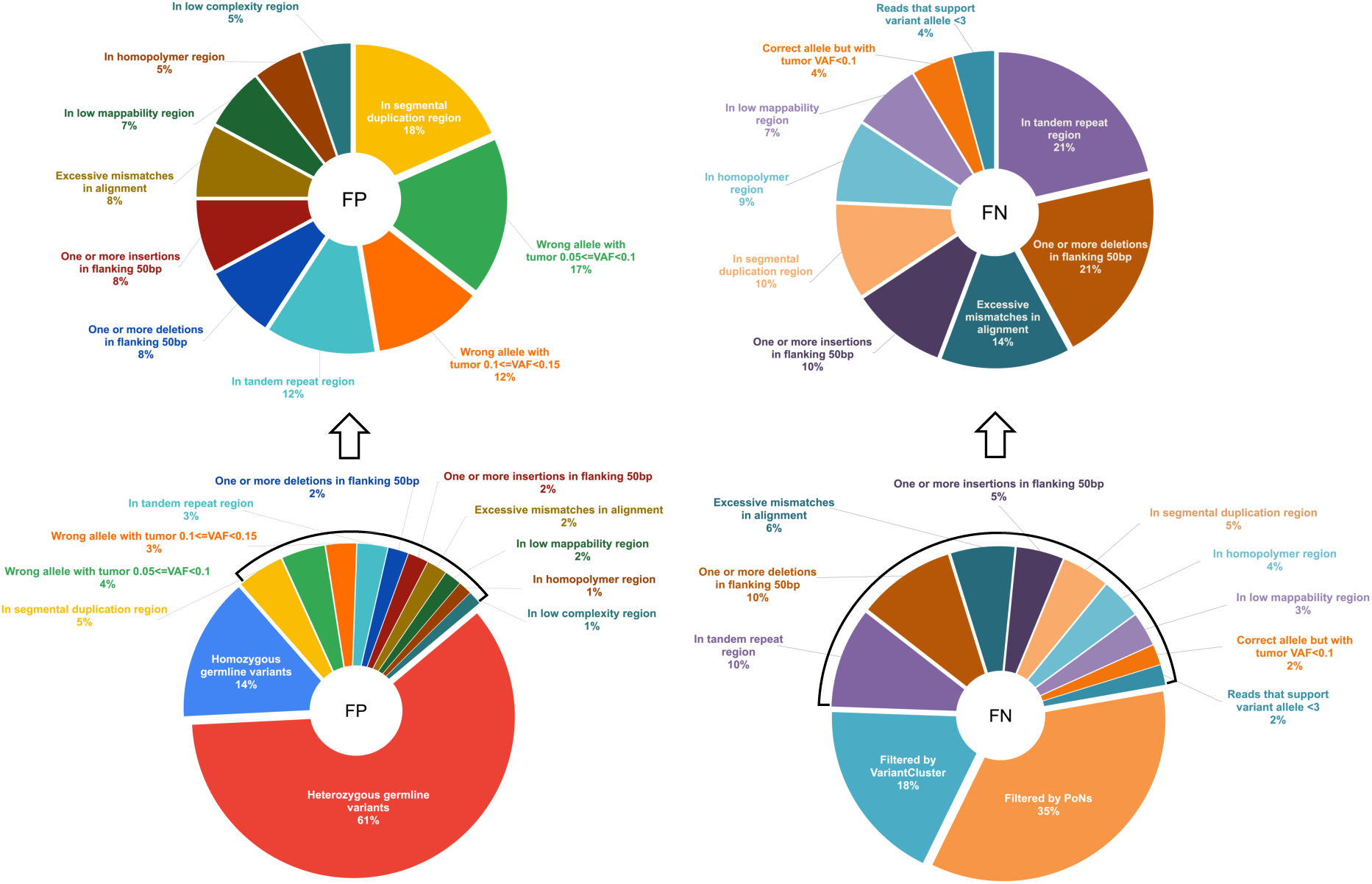
Categorizing the FPs and FNs. Pie charts illustrate the distributions of primary reasons for FPs and FNs in ClairS-TO. A total of 300 FPs and 300 FNs were randomly selected from the calling results on 50-fold coverage of ONT COLO829.Regarding FPs, 61% were heterozygous germline variants, and 14% were homozygous germline variants. The distribution of the remaining FPs (excluding germline variants) is also shown. A major category among these FPs is caused by alignment artifacts in complex genomic contexts, including 18% in segmental duplication regions, 12% in tandem repeat regions, 7% in low-mappability regions, and 5% in homopolymer regions, as defined by GIAB Stratifications v3.3. Regarding FNs, 35% were "Filtered by PoNs" and 18% were "Filtered by VariantCluster", both resulting from post-processing steps. A major category among the remaining FNs (excluding those incorrectly filtered) is also located in complex regions, including 21% in tandem repeat regions, 21% with one or more deletions in the flanking 50 bp, 14% with excessive mismatches in alignment, 10% with one or more insertions in the flanking 50 bp, 10% in segmental duplication regions, 9% in homopolymer regions, and 7% in low-mappability regions.

Regarding FPs, 61% were heterozygous germline variants, and 14% were homozygous germline variants called as germline in Clair3 but as somatic in ClairS-TO. Although most germline variants were tagged by PoNs, the remaining individual germline variants accounted for the majority of FPs. For example, a germline variant with a VAF of 0.6167 at chr3:145,978,865 was called as somatic with a QUAL score of 74.86 in ClairS-TO but as a heterozygous germline with a QUAL score of 28.77 in Clair3. This FP was called due to that ClairS-TO has insufficient evidence to classify it as a germline variant. Among the remaining FPs, 17% and 12% were not reported in the COLO829 truth set, with tumor VAF in the ranges of 0.05≤VAF<0.1 and 0.1≤VAF<0.15, respectively. These variants, some of which are called only with long-read, might have a chance to be correct and be added to the truth set in the future with additional sequencing coverage and orthogonal validations. Another major category of FPs is caused by alignment artifacts in complex genomic contexts, including 18% in segmental duplication regions, 12% in tandem repeat regions, 7% in low-mappability regions, and 5% in homopolymer regions. These issues might be mitigated as sequencing errors in complex regions are reduced.

Regarding FNs, 35% were filtered by PoNs, and 18% were filtered by the "VariantCluster" hard-filter. While these post-filters result in the loss of some true variants, they provide a more substantial improvement in precision. Among the remaining FNs, 8% of the truth variants were missed due to low tumor VAF (VAF<0.1) or insufficient supporting reads (variant allele depth <3), which could potentially be detected at a higher coverage. Other categories of FNs were identified in complex genomic regions, including 21% in tandem repeat regions, 21% with one or more deletions in the flanking 50bp, 14% with excessive mismatches in alignment, 10% with one or more insertions in the flanking 50bp, 10% in segmental duplication regions, 9% in homopolymer regions, and 7% in low-mappability regions.

## Discussion

In this study, we present ClairS-TO, the first publicly available deep-learning-based method for long-read tumor-only somatic variant calling. Our experiments have shown that ClairS-TO is the best-performing caller on ONT and PacBio long-read. When trained with short-reads, ClairS-TO has also shown an edge over well-established short-read tumor-only somatic variant callers. More experiments at multiple coverages, tumor purities, and VAF ranges show that ClairS-TO has maintained the lead. Ablation studies have demonstrated the effectiveness of each component and the solution as a whole.

ClairS-TO uses synthetic tumor samples for model training and can use any number of real tumor samples with verified somatic variant truths for fine-tuning. The practically unlimited number of somatic variants that can be generated from synthesizing tumor samples provide a foundation of using deep-learning for somatic variant calling before more real tumor samples of different cancer types with verified truths are available, considering 1) a model trained only on real tumor samples might risk of overfitting when applied to a cancer type that was not included in training – it is impractical to expect for a model training only on real tumor samples to have a full coverage of all cancer types, and 2) the number of somatic variants in a real sample only ranged from 10k to 100k, which is two orders of magnitude fewer than the germline variants a real sample could offer for model training – it takes more efforts and longer to reach the same number of true variants for training as what we had for deep-learning-based germline variant calling if real tumor sample is the only source. Without a paired normal sample for reference, a tumor-only somatic variant caller is required to not only be capable of extracting variant signals, but also telling somatic variants apart from germline variants and background noises. ClairS-TO uses an ensemble of two neural networks of completely different architectures that were trained from the same samples but for two exact opposite tasks – an affirmative neural network predicts how likely a candidate is a somatic variant and a negational neural network predicts how likely a candidate is not a somatic variant. A false variant that is predicted positive by the affirmative neural network would have a chance to be rejected by a negational prediction from the negational neural network. ClairS-TO uses publicly available and user-supplied PoNs to further remove germline and false variants. Our study found that CoLoRSdb, which was built from long-read sequenced samples, has significantly outperformed 1000G PoN, gnomAD, and dbSNP, which were built primarily based on short-read sequenced samples, in removing non-somatic variants called with long-reads. This urges the community to make public more high-quality long-read sequenced samples for a more comprehensive PoN to further improve the quality of long-read tumor-only somatic variant calling. It also highlights the need to resequence some samples that were used for building in-house PoNs using long-read. The Verdict module that statistically distinguishes somatic variant from germline according to a difference in the expected VAF at different tumor purity, tumor ploidy, and variant copy number profile, was found to be very effective for filtering germline variant, especially when the tumor purity is relatively low, which is often the case of real cancer samples.

Albeit ClairS-TO has outperformed the other callers on all three sequencing platforms, the overall tumor-only Indel calling performance remains low. We believe the algorithm of ClairS-TO remains room to be further improved for a better Indel calling performance, but we highlight the following three tasks that could have a greater potential: 1) A PoN that has a more comprehensive coverage of Indels; 2) Error-correction on Indels using tools such as HERRO^33^ before variant calling; and 3) further advancements in long-read sequencing quality brought by better sequencing chemistries or basecallers.

## Method

### Training data preparation

#### Generating synthetic tumor samples for ClairS-TO SS (synthetic sample) model training

Model training based on synthetic tumor samples was proven effective in the absence of sufficient real tumor samples for model training^5^. The strategy works based on the observation that a germline variant unique to a sample can be considered as a somatic variant when mixed into another sample. A detailed workflow is shown in **Figure 2b**. Using, for example, GIAB HG002 (hereafter referred to as sample A) and GIAB HG001 (B) as sources, the alignments of two samples were split into smaller non-overlapping chunks of about 4-fold coverage each. The small chunks can then be combined to simulate different coverages (through controlling the total number of chunks) and different VAF (through controlling the proportion of chunks from each sample). During the synthesis, three categories of variants, namely "Somatic", "Germline", and "Artifact", can be derived. "Somatic" are those truth germline variants unique to A and not in B, or vice versa. "Germline" are those truth germline variants that appear in both A and B. "Artifact" are those variant candidates in synthetic tumor sample but are not germline variants according to the truths. Only the variants in the intersection of GIAB high-confidence regions of A and B were used for model training. For the "Somatic" and "Germline" variants, we required a minimum coverage of four and a minimum of three reads supporting the variant allele. A summary of different synthetic samples, allelic fraction distributions, and variant categories is shown in **Figure 2c** and **Supplementary Figure 2a**.

#### Using real cancer cell-line samples for ClairS-TO SSRS (synthetic sample and real sample) model training

The generation of synthetic tumor samples can theoretically offer an unlimited number of variants at any tumor purity, normal contamination, or allelic fraction for model training. The synthetic samples are also covering the sequencing error profile well since the reads used for synthesis are real sequencing reads. However, the synthetic samples are unable to cover cancer-specific and cancer-type-specific characteristics, such as mutational signatures^20^. These characteristics only exist in and can only be learned from real tumor samples. Actually, a model agnostic of these characteristics has no problem being used for somatic variant calling, but a model that realizes these characteristics has the potential to achieve better performance.

Park et al.^6^ and Keskus et al.^17^ have released five pairs of real cancer cell-line sample datasets (HCC1395/BL, HCC1937/BL, HCC1954/BL, H1437/BL, and H2009/BL) and used them for model training in DeepSomatic and Severus with success. The real cancer datasets released by the two studies have also enabled ClairS-TO to fine-tune its synthetic samples-based model and achieve further performance improvements.

In addition to excluding chromosome 1 from training, ClairS-TO uses only four real tumor samples (i.e., HCC1937, HCC1954, H1437, and H2009) for model training, and left out the entire HCC1395 for benchmarking purposes. This is unlike Park et al. which has used HCC1395 for model training. To our best knowledge, HCC1395 and COLO829 are so far the two best studied tumor samples with the most confident somatic variants called and reported based on both short-reads and long-reads. HCC1395 particularly, has its truth variants curated by the SEQC2 consortium that involves multiple institutes. However, due to the relatively lower number of somatic variants a real tumor sample usually has, HCC1395 has only 39,560 truth SNVs and 1,922 Indels, and if narrowed down to chromosome 1, only 3,457 SNVs and 147 Indels, leading to insufficient statistical power for reliable performance comparison. In fact, ClairS-TO will benefit from adding HCC1395 to model training, but we have chosen to exclude it to allow future comparisons between ClairS-TO and other works based on the whole HCC1395. On the other hand, HCC1397 and HCC1954 that we have chosen for model training are breast cancer cell lines. Thus, excluding HCC1395, which is also a breast cancer cell line, would be of a lower risk of missing cancer-type-specific characteristics to be learned by ClairS-TO.

"Somatic" and "Germline" variants are provided by Park et al. as VCF files. "Artifact" are those variant candidates in high-confidence regions but were not marked as Somatic or Germline. Similar to synthetic samples, we required a minimum coverage of four and a minimum of three reads supporting the variant allele for Somatic and Germline variants. A summary of different real samples, allelic fraction distributions, and variant categories is shown in **Figure 2d** and **Supplementary Figure 2b**.

### ClairS-TO workflow and design

#### Overview

**Figure 1a** shows an overview of the ClairS-TO somatic variant calling workflow. Given the alignments in the BAM/CRAM format of a tumor sample, ClairS-TO goes through three steps to call the somatic variants and outputs them into a VCF file. In step 1, genome positions with alternative allele supports are chosen as variant candidates, summarized into pileups and converted into tensors, and fed into both the affirmative neural network (AFF) and negational neural network (NEG) to calculate the probabilities of a candidate being an allele as well as not being an allele. In step 2, a posterior probability is computed using the outputs of AFF and NEG, as well as a matrix of prior probabilities derived from the training samples. In step 3, hard-filters, PoNs, and the Verdict module are applied to remove germline and false variants.

#### Step 1: AFF network and NEG network

Principally, every genome position is a variant candidate that can be sent to the neural networks for variant calling. However, very low-coverage positions and those with very low VAF or no alternative allele support are nonstarters that could be excluded. ClairS-TO by default requires a variant candidate to have 1) a minimum coverage of four; 2) three or more alternative allele support reads; and 3) a minimum VAF of 0.05. The alignments at each chosen variant candidate are then summarized into features required by the pileup input. The design of the pileup input is elaborated in **Supplementary Notes** – **Pileup input**. For each variant candidate, the same pileup input is then fed into both the AFF network and NEG network for inference. AFF outputs the probability of a variant candidate being allele "A", "C", "G", or "T". NEG outputs the probability of a variant candidate NOT being allele "A", "C", "G", or "T". When Indel calling is enabled, an additional AFF network and NEG network with the output extended with allele "I" (Insertion) and "D" (Deletion) are used solely for Indel calling. The reasons that we use two separate sets of networks for SNVs and Indels respectively are based on our observation that 1) There exists an order of magnitude more reliable somatic SNVs than somatic Indels for us to train and fine-tune our models – using a single model for calling both SNVs and Indels surprisingly lowered the performance of calling SNVs, and 2) Indel calling is not always required in tumor-only somatic variant calling usage scenarios – using separated models can speed up the calling in some usage scenarios.

#### The choice of network architecture for AFF and NEG

The AFF network uses CvT^34^ (convolutional vision transformer). It is an architecture that incorporates convolution into the transformer. The use of multi-head self-attention and position-wise feed-forward module in CvT makes it excel in modelling the global relations in the input, which makes it suitable for making use of the undiscoverable signals in the flanking alignments of a candidate variant. The aim of introducing the NEG network and using an ensemble of two networks is to reduce the aleatoric uncertainty, thus the architecture of the NEG is supposed to be as different from that of the AFF as possible. We have chosen Bidirectional-GRU as the network architecture of NEG, which is more computationally efficient than but as effective as Bidirectional-LSTM, which was proven to be successful in handling pileup input in Clair3.

#### Step 2: Combining the results of AFF and NEG using Bayesian method

The AFF network outputs *P*_*AFF*_(*y*|*x*) where *x* is the input, and *y* is four possible somatic variant alleles "A", "C", "G", or "T". Similarly, NEG network outputs *P*_*NEG*_(¬*y*|*x*). Ideally, for a variant candidate, *P*_*AFF*_(*y*|*x*) should equal to 1 − *P*_*NEG*_(¬*y*|*x*), that is *P*_*AFF*_(*y*|*x*) and 1 − *P*_*NEG*_(¬*y*|*x*) should be complementary. But it is not always the case because the two probabilities were given by two different models using different network architectures, thus they can make different mistakes. What we expected to see is that in some tricky cases, one network is making a mistake while the other could still make a correct prediction. Therefore, a step is needed to combine the two probabilities. We used Bayesian method to combine the two probabilities, calculated as:

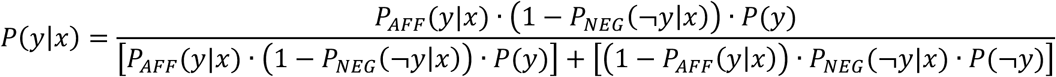

where *P*(*y*) is the prior probability that can be derived from the synthetic tumor samples through counting the proportion of positive training samples at each combination of *P*_*AFF*_(*y*|*x*) and 1 − *P*_*NEG*_(¬*y*|*x*). The distribution of *P*(*y*) is shown in **Supplementary Figure 3**. *P*(¬*y*) equals to 1 − *P*(*y*). The method 1) reinforces agreement – if both models assign high probabilities to the same allele, the combined probability for that allele increases significantly, and 2) mitigates disagreement – if the models disagree, the multiplication reduces the combined probability, reflecting uncertainty.

#### Step 3a. The design of the nine hard-filters to remove false variant calls

1. Exists in two haplotypes: Somatic variants that have a single haplotype origin (either maternal or paternal) should be considered more reliable than those with multiple haplotype origins, except for somatic variants with high VAF that might be a result of copy number alteration or clonal duplication. Input alignments are phased using WhatsHap^35^ or LongPhase^36^. Variants with the alternative allele existing in more than one haplotype are tagged as "MultiHap".
2. No ancestral haplotype support: A somatic variant is considered supported by an ancestral haplotype if the haplotype containing the somatic variant is believed to originate from one of the ancestral haplotypes. Any variants that cannot be found with a supporting ancestral haplotype are tagged as "NoAncestry".
3. Low base quality (BQ): If the average BQ of the alternative alleles <T, the candidate is tagged as "LowAltBQ". T set to 20 as the default.
4. Low mapping quality (MQ): If the average MQ of the alternative alleles <T, the candidate is tagged as "LowAltMQ". T set to 20 as the default.
5. Variant cluster: If three or more variants were called within 200bp, these variants are tagged as "VariantCluster".
6. Read start/end artifacts: Mismatches at the start or end of reads are less reliable due to sequencing and alignment artifacts. If >30% of the supporting alternative alleles of a variant are within the 100bp from the start or end of reads, the candidate is tagged as "ReadStartEnd".
7. Strand bias. If the two-tailed p-value of Fisher Exact Test on the count of forward and reverse strand with alternative allele support against an even distribution is <0.001, a variant is tagged as "StrandBias".
8. Realignment effect (for short-read only): After realignment, if the count of alternative allele supports and VAF both decreased, a variant is tagged as "Realignment".
9. Low sequence entropy: An Indel is more likely to be a false call when the adjacent sequences are of low complexity. For each Indel variant, if the sequence entropy^37^ of the 5-mers of the flanking 32bp from the reference genome is <0.9, the candidate is tagged as "LowSeqEntropy".

#### Step 3b. Panels of normals (PoNs)

As a common practice of tumor-only somatic variant calling^6, 12^, ClairS-TO also uses PoNs to tag non-somatic variants. ClairS-TO by default uses four sources of PoNs, including gnomAD^26^, dbSNP^27^, 1000 genomes project (1000G) PoN^28^, and Consortium of Long Read Sequencing Database (CoLoRSdb)^29^. The original gnomAD r2.1, dbSNP v138, and 1000G PoN databases were downloaded from GATK. After choosing the variants that are with VAF≥0.001 and non-somatic, 35,551,905, 60,683,019, and 2,609,566 have remained for constructing a PoN. The use of CoLoRSdb as a PoN for long-read somatic variant calling was first seen in Park et al. and being introduced into ClairS-TO since version v0.3.0. After choosing those with VAF≥0.001, 49,550,902 variants remained for a PoN. We required exact allele matching for a variant to be tagged as non-somatic for gnomAD and dbSNP, and positional matching for 1000G PoN and CoLoRSdb. Users can also use the "--panel_of_normals" option to make use of additional PoNs and choose whether to require exact allele or positional matching.

#### Step 3c. Verdict, a statistical method to distinguish somatic variant from germline

Verdict in ClairS-TO tags the neural network called variants as either a germline, somatic, or subclonal somatic variant. Verdict’s idea and algorithm are similar to and improved from SGZ^11^. Verdict has three steps, including 1) finding copy number segments, 2) estimating tumor purity, tumor ploidy, and the copy number profile, and 3) binomial tests, as illustrated in **Supplementary Figure 4**. SGZ suggested using ASCAT^38^ to estimate tumor purity, tumor ploidy, and the copy number profile of each variant. We rewrote ASCAT in Python from R so that it could be integrated into Verdict and run reasonably fast. ASCAT uses the LogR (log ratios, representing log-transformed copy numbers derived from sequencing depth) and BAF (B allelic fraction, describing the allelic imbalance of variants) of germline heterozygous variants as input. LogR is calculated by normalizing the read coverage of the tumor sample. BAF is calculated by dividing the signal intensities of minor alleles by those of major and minor alleles. ASCAT uses the LogR and BAF distributions to segment the genome into multiple regions with constant copy number states, identifying breakpoints based on LogR and BAF value changes. Next, ASCAT estimates tumor purity and ploidy by evaluating the goodness of fit for a grid of possible values for tumor purity and ploidy. Using the fact that true copy numbers are nonnegative whole numbers, ASCAT seeks values for tumor purity and ploidy such that the copy number estimates are as close as possible to nonnegative whole numbers for germline heterozygous variants. Then, the allelic fraction of each variant is used as input to two binomial tests that decide whether a variant is more likely to be germline, somatic, or subclonal somatic. The two binomial tests are calculated as follows using the tumor purity, tumor ploidy, and copy number profile as inputs. The two-tailed p-value of the somatic hypothesis (*p*_*somatic*_) is calculated as:

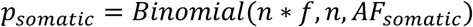

where *n* is the read depth, *f* is the allelic fraction, *AF*_*somatic*_ is the expected allelic fraction of the variant being a somatic, calculated as:

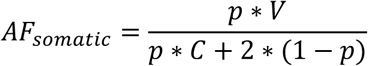

where *p* is the tumor purity, *C* is the copy number, and *V* is the variant allele count in the tumor. The p-value of the germline hypothesis (*p*_*germline*_) is calculated as:

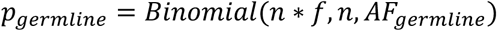

where *AF*_*germline*_ is the expected allelic fraction of the variant being a germline, calculated as:

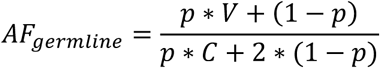

Verdict tags a variant as 1) somatic if *p*_*somatic*_ is greater than 0.001 and *p*_*germline*_ is lower than 0.001; 2) germline if *p*_*somatic*_ is lower than 0.001 and *p*_*germline*_ is greater than 0.001; or 3) subclonal somatic if both *p*_*somatic*_ and *p*_*germline*_ are lower than 0.001, tumor purity is greater than 0.2, and *f* < ^*AFsomatic*^/_*γ*_, where *γ* is a tunable parameter with a default of 1.5.

#### Output

ClairS-TO supports VCF format output. A variant call is tagged as "LowQual" if the variant quality is low (i.e., QUAL<4, configurable by option); "NonSomatic" if exists in PoNs; "MultiHap", "NoAncestry", "LowAltBQ", "LowAltMQ", "VariantCluster", "ReadStartEnd", "StrandBias", "Realignment", or "LowSeqEntropy" if not passing a corresponding hard-filter; and "Verdict_Germline", "Verdict_Somatic", or "Verdict_SubclonalSomatic" as predicted by the Verdict module. For each variant, the support information such as allelic fraction, read depth, allele depth of both strands are given in the VCF file.

### Benchmarking

We used two cancer cell lines COLO829 and HCC1395 for benchmarking. The truth somatic variants of COLO829 are provided by NYGC^21^, with a total of 42,993 SNVs and 985 Indels. The truths of HCC1395 are provided by the SEQC2 consortium^22^. We only used those labeled as "HighConf" (high confidence) and "MedConf" (medium confidence) in the high-confidence regions, with a total of 39,447 SNVs and 1,602 Indels. To reflect the real performance of the callers and remove the false negative calls that are actually caused by no coverage or no alternative allele support at all in the input, we required a truth to be included for benchmarking if it has 1) coverage ≥4; 2) reads supporting an alternative allele ≥3; and 3) VAF ≥0.05. We used "som.py" in Illumina’s Haplotype Comparison Tools^39^ to generate precision, recall, and F1-score metrics. The "compare_vcf" submodule in ClairS-TO generates the same metrics and has automated the exclusion of unqualified truth variants. We cross-validated the metrics generated by both "som.py" and "compare_vcf". The command lines we used to run each caller and benchmarks are given in the **Supplementary Notes** – **Command lines used** section.

### Computational performance

ClairS-TO was implemented primarily with Python and PyTorch, with performance-critical parts written in C++ or accelerated with PyPy. Model training of ClairS-TO requires a high-end graphics processing unit (GPU). In this study, we used Nvidia GeForce RTX 3090 and 4090 for model training. For variant calling, ClairS-TO requires only CPU resources. Using two 12-core Intel Xeon Silver 4116 processors, ClairS-TO finished variant calling on 50-fold coverage of ONT Q20+ COLO829 WGS data in 214 minutes, using 1GB of memory per process at peak. In comparison, DeepSomatic used 516 minutes. For 50-fold coverage of Illumina COLO829 WGS data, ClairS-TO, Mutect2, Octopus, and Pisces took 211, 971, 3,211, and 1,785 minutes, respectively.

## Code availability

ClairS-TO is open source and available at https://github.com/HKU-BAL/ClairS-TO under the BSD 3-Clause license. The results in this paper were based on the ClairS-TO latest release (version 0.4.0). Multiple installation options are available for ClairS-TO, including Docker, Singularity, and Conda.

## Data availability

All data used in this study, including the 1) ONT, PacBio, and Illumina sequencing datasets; 2) GIAB truth variants, confident regions, and stratifications; and 3) reference genomes are publicly accessible via links or SRA accession IDs listed in the **Supplementary Notes** – **Data availability** section. A summary of the sequencing data used for model training and benchmarking is shown in **Supplementary Table 1**. All analysis output, including the VCFs, are available at http://www.bio8.cs.hku.hk/clairs-to/analysis_result.

## Acknowledgements

R.L. was supported by Hong Kong Research Grants Council grants GRF (17113721) and CRF (C7003-24Y), the URC fund at HKU, and Oxford Nanopore Technologies.

## Author contributions

R. L. conceived the study. L. C., Z. Z., and R. L. designed the algorithms, implemented and benchmarked ClairS-TO, and wrote the paper. J. S., X. Y., A. O. K. W., J. Z. and Y. L. evaluated the benchmarking results. All authors revised the manuscript.

