## Supplementary Materials for "ClairS-TO: A deep-learning method for long-read tumor-only somatic small variant calling"

### Supplementary Notes

Running other somatic variant callers for Illumina data ... 16
*DeepSomatic (v1.7.0)* ... 16
*Mutect2 (v4.2.6.1)* ... 17
*Octopus (v0.7.4)* ... 17
*Pisces (v5.3.0)* ... 17
Benchmarking ... 17
*som.py (v0.3.12)* ... 17
*Calculate Precision, Recall, F1-Score with "compare_vcf" submodule in ClairS-TO* ... 18

**DATA AVAILABILITY ... 18**

GIAB truth variants ... 18
*HG001 (NA12878), GRCh38, v4.2.1* ... 18
*HG002 (NA24385), GRCh38, v4.2.1* ... 18
*HG003 (NA24149), GRCh38, v4.2.1* ... 18
*HG004 (NA24143), GRCh38, v4.2.1* ... 18
NYGC truth variants for COLO829 sample ... 19
*COLO829, GRCh38* ... 19
SEQC truth variants for HCC1395 sample ... 19
*HCC1395, GRCh38, v1.2* ... 19
Four real cancer cell-lines variants from Park et al. ... 19
*HCC1937, GRCh38* ... 19
*HCC1954, GRCh38* ... 19
*H1437, GRCh38* ... 19
*H2009, GRCh38* ... 19
Reference genomes ... 20
*GRCh38_no_alt* ... 20
*GRCh38 d1.vd1* ... 20
*GRCh38 Stratification regions (v3.3)* ... 20
ONT Sequencing Data ... 20
*ONT EPI2ME Labs HG001 R10.4.1 Q20+, GRCh38_no_alt, 78.89-fold* ... 20
*ONT EPI2ME Labs HG002 R10.4.1 Q20+, GRCh38_no_alt, 91.18-fold* ... 20
*Park et al. HCC1937 R10.4.1 Q20+, GRCh38_no_alt, 158.14-fold* ... 20
*Park et al. HCC1954 R10.4.1 Q20+, GRCh38_no_alt, 82.44-fold* ... 21
*Park et al. H1437 R10.4.1 Q20+, GRCh38_no_alt, 79.10-fold* ... 21
*Park et al. H2009 R10.4.1 Q20+, GRCh38_no_alt, 106.92-fold* ... 21
*Park et al. HCC1395 R10.4.1 Q20+, GRCh38_no_alt, 79.52-fold* ... 21
*ONT EPI2ME Labs COLO829 R10.4.1 Q20+, GRCh38_no_alt, 101.80-fold* ... 21
PacBio Sequencing Data ... 21
*PacBio HG003 HiFi Revio, GRCh38_no_alt, 61.15-fold* ... 21
*PacBio HG004 HiFi Revio, GRCh38_no_alt, 62.67-fold* ... 21
*Park et al. HCC1937 HiFi Revio, GRCh38_no_alt, 59.50-fold* ... 22
*Park et al. HCC1954 HiFi Revio, GRCh38_no_alt, 63.07-fold* ... 22
*Park et al. H1437 HiFi Revio, GRCh38_no_alt, 64.06-fold* ... 22
*Park et al. H2009 HiFi Revio, GRCh38_no_alt, 64.89-fold* ... 22
*PacBio COLO829 HiFi Revio, GRCh38_no_alt, 60.70-fold* ... 22
Illumina Sequencing Data ... 22
*HG003 NovaSeq 6000 (NA24149), GRCh38_no_alt, 47.38-fold* ... 22
*HG004 NovaSeq 6000 (NA24385), GRCh38_no_alt, 46.36-fold* ... 22
*HG003 HiSeqX (NA24149), GRCh38_no_alt 43.77-fold* ... 23
*HG004 HiSeqX (NA24385), GRCh38_no_alt, 42.13-fold* ... 23
*Park et al. HCC1937 NovaSeq 6000, GRCh38, 256.28-fold* ... 23

### Supplementary Figures

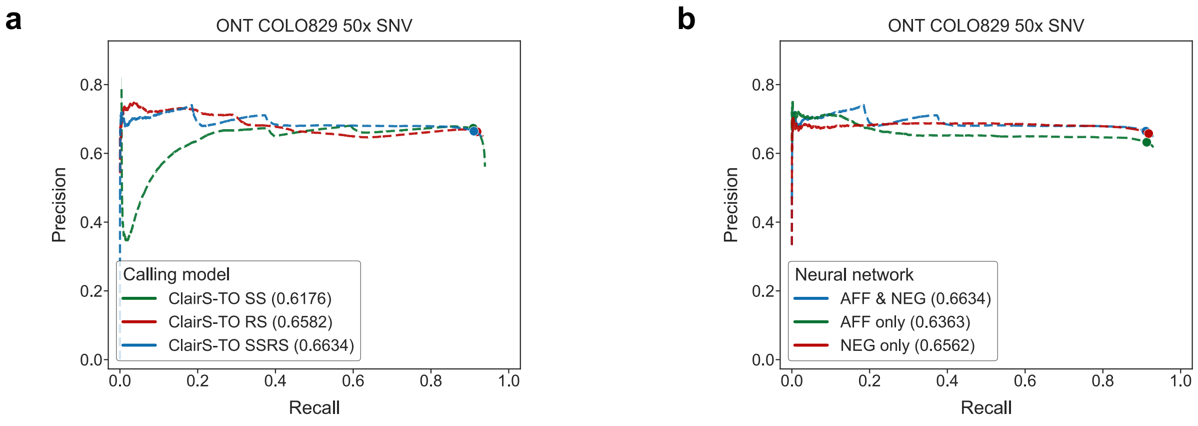

**Supplementary Figure 1. Performance evaluation with different model settings**

(a) The Precision-Recall curves on 50-fold coverage of ONT COLO829 of different ClairS-TO models: ClairS-TO SS, ClairS-TO RS, and ClairS-TO SSRS. The dot on each dashed line represents the best F1-score achieved. AUPRC is shown in parentheses in the legend. The results show that ClairS-TO SSRS outperformed the other two models in terms of AUPRC for both SNV and Indel. (b) The Precision-Recall curves on 50-fold coverage of ONT COLO829 using 1) both AFF & NEG, 2) only AFF, or 3) only NEG in the SSRS model. The dot on each dashed line represents the best F1-score achieved. AUPRC is shown in curve brackets in the legend.

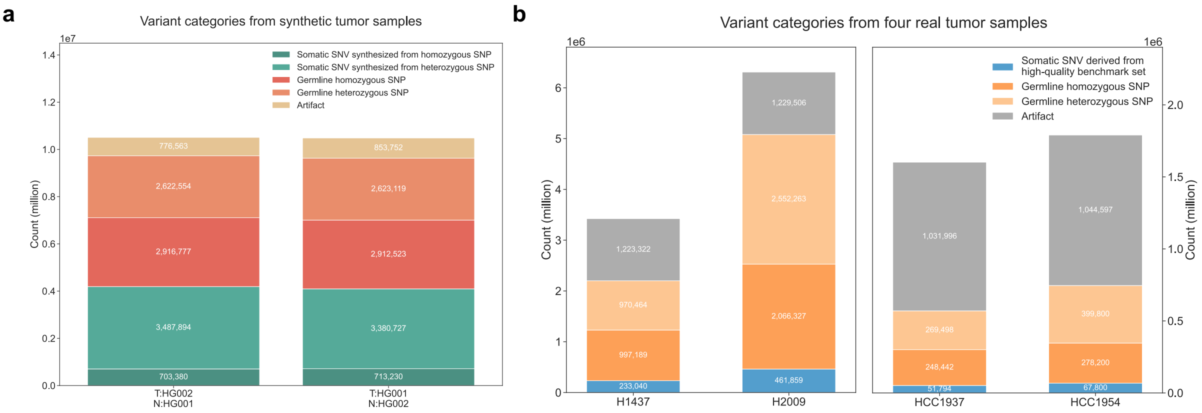

**Supplementary Figure 2. Number summary of synthetic/real tumor samples and variant categories**

(a) The breakdown of the number of "Somatic", "Germline", and "Artifact" variants in the two synthetic tumor samples used for model training. (b) The breakdown of the number of "Somatic", "Germline", and "Artifact" variants in the four real tumor samples used for model training.

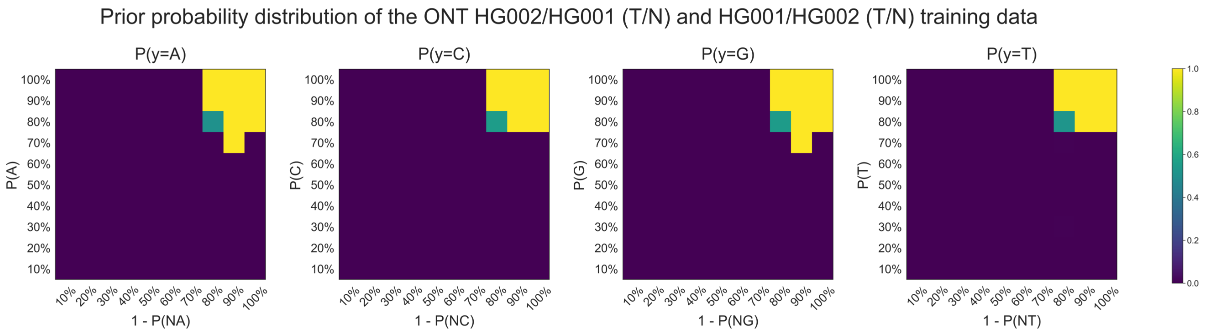

**Supplementary Figure 3. Prior probability distribution visualization**

The figure shows the prior probability distribution generated from ONT HG002/HG001 (T/N) and HG001/HG002 (T/N) training data. The prior probabilities were generated with the following steps. First, for each probability output in both the AFF ($P(A)$, $P(C)$, $P(G)$, and $P(T)$) and NEG ($P(NA)$, $P(NC)$, $P(NG)$, and $P(NT)$) networks, the values of all training candidates are ranked and derived into 10 bins. Next, four 10 by 10 matrices (i.e., $P(y = A)$, $P(y = C)$, $P(y = G)$, and $P(y = T)$)) are obtained according to the truths (how many training candidates are positive in each bin). The four matrices serve as the prior probabilities (i.e., $P(y)$) for calculating the joint posterior probability of a candidate being a somatic variant (i.e., $P(y|x)$).

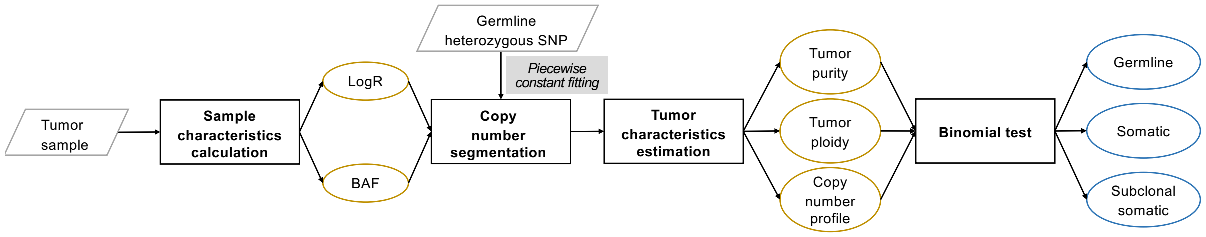

**Supplementary Figure 4. Overview of Verdict workflow**

The figure shows an overview of the workflow of the Verdict module used for post-processing. Verdict distinguishes somatic variants from germlines through three steps: 1) identify copy number segments, 2) estimate tumor sample purity, tumor sample ploidy, and the copy number for each variant candidate, and 3) perform binomial tests for three possible categories including "Germline", "Somatic", and "Subclonal somatic", and choose the category with the highest probability.

### Supplementary Tables

**Supplementary Table 1. Summary of datasets for model training and performance evaluation**

(a) A summary of sequencing data in ONT, PacBio, and Illumina used for model training and benchmarking. (b) The number of different categories of variants in the two synthetic tumor samples (HG002/HG001 (T/N), HG001/HG002 (T/N)). (c) The number of different categories of variants in the four real cancer cell line samples (HCC1937, HCC1954, H1437, and H2009).

**a**

| Platform | Sample | Reference | Aligner | Coverage | Source | Chemistry /Instrument | Used in training | Used for benchmarking |
|---|---|---|---|---|---|---|---|---|
| ONT | HG001 | GRCh38 | Minimap2 | 78.89 | ONT EPI2ME Labs | R10.4.1 | ✓ | |
| | HG002 | GRCh38 | Minimap2 | 91.18 | ONT EPI2ME Labs | R10.4.1 | ✓ | |
| | HCC1937 | GRCh38 | Minimap2 | 158.14 | UCSC and Google | R10.4.1 | ✓ | |
| | HCC1954 | GRCh38 | Minimap2 | 82.44 | UCSC and Google | R10.4.1 | ✓ | |
| | H1437 | GRCh38 | Minimap2 | 79.10 | UCSC and Google | R10.4.1 | ✓ | |
| | H2009 | GRCh38 | Minimap2 | 106.92 | UCSC and Google | R10.4.1 | ✓ | |
| | HCC1395 | GRCh38 | Minimap2 | 79.52 | UCSC and Google | R10.4.1 | | ✓ |
| | COLO829 | GRCh38 | Minimap2 | 101.80 | ONT EPI2ME Labs | R10.4.1 | | ✓ |
| PacBio | HG003 | GRCh38 | pbmm2 | 61.15 | PacBio | HiFi Revio | ✓ | |
| | HG004 | GRCh38 | pbmm2 | 62.67 | PacBio | HiFi Revio | ✓ | |
| | HCC1937 | GRCh38 | pbmm2 | 59.50 | UCSC and Google | HiFi Revio | ✓ | |
| | HCC1954 | GRCh38 | pbmm2 | 63.07 | UCSC and Google | HiFi Revio | ✓ | |
| | H1437 | GRCh38 | pbmm2 | 64.06 | UCSC and Google | HiFi Revio | ✓ | |
| | H2009 | GRCh38 | pbmm2 | 64.89 | UCSC and Google | HiFi Revio | ✓ | |
| | COLO829 | GRCh38 | pbmm2 | 60.70 | PacBio | HiFi Revio | | ✓ |
| Illumina | HG003 | GRCh38 | BWA-MEM | 47.38 | Google Health Center | NovaSeq 6000 | ✓ | |
| | HG004 | GRCh38 | BWA-MEM | 46.36 | Google Health Center | NovaSeq 6000 | ✓ | |
| | HG003 | GRCh38 | BWA-MEM | 43.77 | Google Health Center | HiseqX | ✓ | |
| | HG004 | GRCh38 | BWA-MEM | 42.13 | Google Health Center | HiseqX | ✓ | |
| | HCC1937 | GRCh38 | BWA-MEM | 256.28 | UCSC and Google | NovaSeq 6000 | ✓ | |
| | HCC1954 | GRCh38 | BWA-MEM | 139.02 | UCSC and Google | NovaSeq 6000 | ✓ | |
| | H1437 | GRCh38 | BWA-MEM | 203.94 | UCSC and Google | NovaSeq 6000 | ✓ | |
| | H2009 | GRCh38 | BWA-MEM | 230.57 | UCSC and Google | NovaSeq 6000 | ✓ | |
| | COLO829 | GRCh38 | BWA-MEM | 234.44 | dbGaP | NovaSeq 6000 | | ✓ |

**b**

| Type | Category | Synthetic tumor samples | | |
|---|---|---|---|---|
| | | HG002/HG001 (T/N) | HG001/HG002 (T/N) | Total |
| SNV | Somatic SNV synthesized from homozygous SNP | 703,380 | 713,230 | 1,416,610 |
| | Somatic SNV synthesized from heterozygous SNP | 3,487,894 | 3,380,727 | 6,868,621 |
| | Germline homozygous SNP | 2,916,777 | 2,912,523 | 5,829,300 |
| | Germline heterozygous SNP | 2,622,554 | 2,623,119 | 5,245,673 |
| | Artifact | 776,563 | 853,752 | 1,630,315 |
| Indel | Somatic Indel synthesized from homozygous Indel | 66,134 | 64,129 | 130,263 |
| | Somatic Indel synthesized from heterozygous Indel | 324,426 | 298,573 | 622,999 |
| | Germline homozygous Indel | 265,847 | 265,853 | 531,700 |
| | Germline heterozygous Indel | 232,698 | 232,774 | 465,472 |
| | Artifact | 6,367,467 | 7,355,475 | 13,722,942 |

**c**

| Type | Category | Real tumor samples | | | | |
|---|---|---|---|---|---|---|
| | | HCC1937 | HCC1954 | H1437 | H2009 | Total |
| SNV | Somatic SNV derived from high-quality benchmark set | 51,794 | 67,800 | 233,040 | 461,859 | 814,493 |
| | Germline homozygous SNP | 248,442 | 278,200 | 997,189 | 2,066,327 | 3,590,158 |
| | Germline heterozygous SNP | 269,498 | 399,800 | 970,464 | 2,552,263 | 4,192,025 |
| | Artifact | 1,031,996 | 1,044,597 | 1,223,322 | 1,229,506 | 4,529,421 |
| Indel | Somatic Indel derived from high-quality benchmark set | 3,412 | 10,463 | 14,973 | 17,777 | 46,625 |
| | Germline homozygous Indel | 13,368 | 37,864 | 69,986 | 68,675 | 189,893 |
| | Germline heterozygous Indel | 20,752 | 66,766 | 79,744 | 109,095 | 276,357 |
| | Artifact | 68,240 | 209,260 | 299,460 | 355,540 | 932,500 |

#### Supplementary Table 2. Performance on ONT COLO829 and HCC1395 datasets at different sequencing coverages

(a) The results of ClairS-TO (SS and SSRS model), Clair3, and DeepSomatic for SNV and Indel at ONT COLO829 25-, 50-, and 75-fold tumor coverages. (b) The results of ClairS-TO (SS and SSRS model) and Clair3 for SNV and Indel at ONT HCC1395 25-, 50-, and 75-fold tumor coverages. DeepSomatic was excluded when benchmarking on HCC1395 as DeepSomatic included HCC1395 in model training.

**a**

| Dataset | Caller | Tumor coverage | Type | AUPRC | Precision | Recall | F1-score | TP | FP | FN |
|---|---|---|---|---|---|---|---|---|---|---|
| COLO829 | ClairS-TO SS | 75x | SNV | 0.6191 | 0.6475 | 0.9275 | 0.7626 | 34,810 | 18,951 | 2,723 |
| | | | Indel | 0.1862 | 0.1961 | 0.4872 | 0.2797 | 285 | 1,168 | 300 |
| | | 50x | SNV | 0.6176 | 0.6486 | 0.9282 | 0.7636 | 34,837 | 18,870 | 2,696 |
| | | | Indel | 0.1833 | 0.1963 | 0.4889 | 0.2801 | 286 | 1,171 | 299 |
| | | 25x | SNV | 0.6153 | 0.6556 | 0.9135 | 0.7633 | 34,288 | 18,015 | 3,245 |
| | | | Indel | 0.1665 | 0.1849 | 0.4786 | 0.2668 | 280 | 1,234 | 305 |
| | ClairS-TO SSRS | 75x | SNV | 0.6685 | 0.6606 | 0.9155 | 0.7675 | 34,362 | 17,651 | 3,171 |
| | | | Indel | 0.2062 | 0.2032 | 0.5179 | 0.2919 | 303 | 1,188 | 282 |
| | | 50x | SNV | 0.6634 | 0.6614 | 0.9163 | 0.7683 | 34,391 | 17,605 | 3,142 |
| | | | Indel | 0.2019 | 0.1974 | 0.4957 | 0.2824 | 290 | 1,179 | 295 |
| | | 25x | SNV | 0.6489 | 0.6643 | 0.9061 | 0.7666 | 34,010 | 17,187 | 3,523 |
| | | | Indel | 0.1870 | 0.1926 | 0.4564 | 0.2709 | 267 | 1,119 | 318 |
| | Clair3 | 75x | SNV | 0.5587 | 0.6202 | 0.8112 | 0.7029 | 30,446 | 18,647 | 7,087 |
| | | | Indel | 0.1064 | 0.1397 | 0.3299 | 0.1962 | 193 | 1,189 | 392 |
| | | 50x | SNV | 0.5686 | 0.6239 | 0.8310 | 0.7127 | 31,191 | 18,804 | 6,342 |
| | | | Indel | 0.1040 | 0.1300 | 0.3624 | 0.1913 | 212 | 1,419 | 373 |
| | | 25x | SNV | 0.5543 | 0.5893 | 0.8120 | 0.6830 | 30,477 | 21,236 | 7,056 |
| | | | Indel | 0.0675 | 0.0882 | 0.3573 | 0.1415 | 209 | 2,161 | 376 |
| | DeepSomatic | 75x | SNV | 0.6365 | 0.5952 | 0.9059 | 0.7184 | 34,002 | 23,124 | 3,531 |
| | | | Indel | 0.1190 | 0.1113 | 0.1385 | 0.1234 | 81 | 647 | 504 |
| | | 50x | SNV | 0.6334 | 0.5993 | 0.9157 | 0.7245 | 34,368 | 22,978 | 3,165 |
| | | | Indel | 0.0992 | 0.1030 | 0.1350 | 0.1169 | 79 | 688 | 506 |
| | | 25x | SNV | 0.6030 | 0.5853 | 0.8794 | 0.7028 | 33,005 | 23,387 | 4,528 |
| | | | Indel | 0.0644 | 0.0597 | 0.2085 | 0.0929 | 122 | 1,920 | 463 |

**b**

| Dataset | Caller | Tumor coverage | Type | AUPRC | Precision | Recall | F1-score | TP | FP | FN |
|---|---|---|---|---|---|---|---|---|---|---|
| HCC1395 | ClairS-TO SS | 75x | SNV | 0.5902 | 0.7013 | 0.9231 | 0.7971 | 29,734 | 12,663 | 2,478 |
| | | | Indel | 0.4340 | 0.4379 | 0.6807 | 0.5329 | 923 | 1,185 | 433 |
| | | 50x | SNV | 0.5855 | 0.7047 | 0.9066 | 0.7930 | 29,204 | 12,239 | 3,008 |
| | | | Indel | 0.4159 | 0.4185 | 0.6586 | 0.5117 | 893 | 1,241 | 463 |
| | | 25x | SNV | 0.5690 | 0.6897 | 0.8537 | 0.7630 | 27,499 | 12,371 | 4,713 |
| | | | Indel | 0.4072 | 0.5253 | 0.3909 | 0.4482 | 530 | 479 | 826 |
| | ClairS-TO SSRS | 75x | SNV | 0.6012 | 0.7150 | 0.9047 | 0.7987 | 29,141 | 11,617 | 3,071 |
| | | | Indel | 0.4396 | 0.5169 | 0.5642 | 0.5395 | 765 | 715 | 591 |
| | | 50x | SNV | 0.5986 | 0.7128 | 0.8970 | 0.7943 | 28,893 | 11,643 | 3,319 |
| | | | Indel | 0.4267 | 0.4968 | 0.5206 | 0.5085 | 706 | 715 | 650 |
| | | 25x | SNV | 0.5788 | 0.7038 | 0.8441 | 0.7676 | 27,189 | 11,444 | 5,023 |
| | | | Indel | 0.3996 | 0.4715 | 0.4204 | 0.4444 | 570 | 639 | 786 |
| | Clair3 | 75x | SNV | 0.4817 | 0.6073 | 0.6635 | 0.6341 | 21,374 | 13,824 | 10,838 |
| | | | Indel | 0.2915 | 0.3097 | 0.4661 | 0.3721 | 632 | 1,409 | 724 |
| | | 50x | SNV | 0.4798 | 0.5952 | 0.6722 | 0.6314 | 21,652 | 14,723 | 10,560 |
| | | | Indel | 0.2893 | 0.3156 | 0.4373 | 0.3666 | 593 | 1,286 | 763 |
| | | 25x | SNV | 0.4296 | 0.5743 | 0.6146 | 0.5938 | 19,798 | 14,673 | 12,414 |
| | | | Indel | 0.2245 | 0.2801 | 0.3864 | 0.3248 | 524 | 1,347 | 832 |

#### Supplementary Table 3. Performance on ONT COLO829 dataset at different VAF ranges

The results of ClairS-TO (SS and SSRS model), Clair3, and DeepSomatic for SNV and Indel on 50-fold coverage of ONT COLO829 at different VAF ranges.

| Dataset | Caller | Tumor coverage | AF range | Type | Precision | Recall | F1-score | TP | FP | FN |
|---|---|---|---|---|---|---|---|---|---|---|
| COLO829 | ClairS-TO SS | 50x | 0.05-0.2 | SNV | 0.1631 | 0.6908 | 0.2638 | 680 | 3,131 | 273 |
| | | | | Indel | 0.0592 | 0.1149 | 0.0781 | 14 | 159 | 77 |
| | | | 0.2-0.3 | SNV | 0.6346 | 0.9148 | 0.7494 | 2,675 | 1,503 | 243 |
| | | | | Indel | 0.2318 | 0.4070 | 0.2954 | 45 | 116 | 51 |
| | | | 0.3-0.4 | SNV | 0.6954 | 0.9359 | 0.7979 | 4,908 | 2,123 | 332 |
| | | | | Indel | 0.2018 | 0.4314 | 0.2750 | 58 | 174 | 58 |
| | | | 0.4-0.5 | SNV | 0.6996 | 0.9526 | 0.8067 | 7,265 | 3,106 | 360 |
| | | | | Indel | 0.2045 | 0.7273 | 0.3192 | 68 | 249 | 24 |
| | | | 0.5-1.0 | SNV | 0.6595 | 0.9375 | 0.7743 | 19,707 | 10,159 | 1,311 |
| | | | | Indel | 0.1642 | 0.7209 | 0.2675 | 159 | 789 | 60 |
| | ClairS-TO SSRS | 50x | 0.05-0.2 | SNV | 0.2330 | 0.5572 | 0.3285 | 569 | 1,620 | 391 |
| | | | | Indel | 0.1667 | 0.0115 | 0.0215 | 1 | 5 | 86 |
| | | | 0.2-0.3 | SNV | 0.6615 | 0.8752 | 0.7535 | 2,591 | 1,278 | 356 |
| | | | | Indel | 0.3214 | 0.3140 | 0.3176 | 27 | 57 | 59 |
| | | | 0.3-0.4 | SNV | 0.7081 | 0.9102 | 0.7966 | 4,828 | 1,943 | 465 |
| | | | | Indel | 0.2143 | 0.5000 | 0.3000 | 52 | 187 | 51 |
| | | | 0.4-0.5 | SNV | 0.7085 | 0.9428 | 0.8090 | 7,224 | 2,946 | 434 |
| | | | | Indel | 0.1959 | 0.7614 | 0.3116 | 67 | 275 | 21 |
| | | | 0.5-1.0 | SNV | 0.6655 | 0.9306 | 0.7760 | 19,675 | 9,818 | 1,456 |
| | | | | Indel | 0.1683 | 0.7070 | 0.2719 | 154 | 751 | 63 |
| | Clair3 | 50x | 0.05-0.2 | SNV | 0.0000 | 0.0000 | 0.0000 | 62 | 9 | 883 |
| | | | | Indel | 0.0000 | 0.0000 | 0.0000 | 8 | 2 | 87 |
| | | | 0.2-0.3 | SNV | 0.6337 | 0.1661 | 0.2633 | 1,915 | 274 | 2,379 |
| | | | | Indel | 0.1364 | 0.0349 | 0.0556 | 46 | 19 | 83 |
| | | | 0.3-0.4 | SNV | 0.6413 | 0.7432 | 0.6885 | 4,873 | 2,153 | 1,330 |
| | | | | Indel | 0.1138 | 0.1863 | 0.1413 | 78 | 148 | 83 |
| | | | 0.4-0.5 | SNV | 0.6383 | 0.9546 | 0.7650 | 7,439 | 4,108 | 345 |
| | | | | Indel | 0.1506 | 0.6591 | 0.2452 | 83 | 327 | 30 |
| | | | 0.5-1.0 | SNV | 0.6154 | 0.9350 | 0.7423 | 20,562 | 12,260 | 1,365 |
| | | | | Indel | 0.1251 | 0.6140 | 0.2079 | 198 | 923 | 83 |
| | DeepSomatic | 50x | 0.05-0.2 | SNV | 0.2535 | 0.3092 | 0.2786 | 708 | 804 | 610 |
| | | | | Indel | 0.1667 | 0.0115 | 0.0215 | 8 | 5 | 86 |
| | | | 0.2-0.3 | SNV | 0.5881 | 0.7767 | 0.6694 | 2,731 | 1,552 | 637 |
| | | | | Indel | 0.1974 | 0.1744 | 0.1852 | 30 | 61 | 71 |
| | | | 0.3-0.4 | SNV | 0.6353 | 0.9004 | 0.7449 | 5,050 | 2,677 | 516 |
| | | | | Indel | 0.1165 | 0.1176 | 0.1171 | 31 | 91 | 90 |
| | | | 0.4-0.5 | SNV | 0.6351 | 0.9424 | 0.7588 | 7,457 | 4,112 | 437 |
| | | | | Indel | 0.1209 | 0.2500 | 0.1630 | 34 | 160 | 66 |
| | | | 0.5-1.0 | SNV | 0.5919 | 0.9559 | 0.7311 | 20,514 | 13,833 | 925 |
| | | | | Indel | 0.0725 | 0.1349 | 0.0943 | 82 | 371 | 186 |

**Supplementary Table 4. Performance on ONT COLO829 dataset at different tumor purities**

(a) The results of ClairS-TO (SS and SSRS model), Clair3, and DeepSomatic for SNV and Indel on 50-fold coverage of ONT COLO829 at five tumor purities. (b) The performance of ClairS-TO (SS and SSRS model) on 50-fold coverage of ONT COLO829 with or without Verdict module.

a

| Dataset | Caller | Tumor coverage | Tumor purity | Type | AUPRC | Precision | Recall | F1-score | TP | FP | FN |
|---|---|---|---|---|---|---|---|---|---|---|---|
| COLO829 | ClairS-TO SS | 50x | 1.0 | SNV | 0.6176 | 0.6486 | 0.9282 | 0.7636 | 34,837 | 18,870 | 2,696 |
| | | | | Indel | 0.1833 | 0.1963 | 0.4889 | 0.2801 | 286 | 1,171 | 299 |
| | | | 0.8 | SNV | 0.5930 | 0.6389 | 0.9118 | 0.7513 | 34,223 | 19,343 | 3,310 |
| | | | | Indel | 0.1637 | 0.1712 | 0.4838 | 0.2529 | 283 | 1,370 | 302 |
| | | | 0.6 | SNV | 0.5849 | 0.6642 | 0.8579 | 0.7488 | 32,200 | 16,277 | 5,333 |
| | | | | Indel | 0.1695 | 0.1990 | 0.3350 | 0.2497 | 196 | 789 | 389 |
| | | | 0.4 | SNV | 0.5470 | 0.6845 | 0.8242 | 0.7479 | 30,936 | 14,261 | 6,597 |
| | | | | Indel | 0.1481 | 0.1349 | 0.4205 | 0.2042 | 246 | 1,578 | 339 |
| | | | 0.2 | SNV | 0.4851 | 0.6318 | 0.7102 | 0.6687 | 26,655 | 15,532 | 10,878 |
| | | | | Indel | 0.1223 | 0.0596 | 0.3248 | 0.1007 | 190 | 2,999 | 395 |
| | ClairS-TO SSRS | 50x | 1.0 | SNV | 0.6634 | 0.6614 | 0.9163 | 0.7683 | 34,391 | 17,605 | 3,142 |
| | | | | Indel | 0.2019 | 0.1974 | 0.4957 | 0.2824 | 290 | 1,179 | 295 |
| | | | 0.8 | SNV | 0.6256 | 0.6637 | 0.8852 | 0.7586 | 33,224 | 16,832 | 4,309 |
| | | | | Indel | 0.1767 | 0.1889 | 0.4171 | 0.2600 | 244 | 1,048 | 341 |
| | | | 0.6 | SNV | 0.6041 | 0.6695 | 0.8560 | 0.7514 | 32,128 | 15,857 | 5,405 |
| | | | | Indel | 0.1697 | 0.1942 | 0.3573 | 0.2517 | 209 | 867 | 376 |
| | | | 0.4 | SNV | 0.5500 | 0.6862 | 0.8255 | 0.7495 | 30,984 | 14,166 | 6,549 |
| | | | | Indel | 0.1243 | 0.1552 | 0.2462 | 0.1904 | 144 | 784 | 441 |
| | | | 0.2 | SNV | 0.4797 | 0.6992 | 0.6430 | 0.6699 | 24,132 | 10,384 | 13,401 |
| | | | | Indel | 0.0788 | 0.0890 | 0.0855 | 0.0872 | 50 | 512 | 535 |
| | Clair3 | 50x | 1.0 | SNV | 0.5686 | 0.6239 | 0.8310 | 0.7127 | 31,191 | 18,804 | 6,342 |
| | | | | Indel | 0.1040 | 0.1300 | 0.3624 | 0.1913 | 212 | 1,419 | 373 |
| | | | 0.8 | SNV | 0.3378 | 0.4991 | 0.6411 | 0.5613 | 24,064 | 24,154 | 13,469 |
| | | | | Indel | 0.0515 | 0.0656 | 0.3573 | 0.1109 | 209 | 2,975 | 376 |
| | | | 0.6 | SNV | 0.1494 | 0.2934 | 0.3769 | 0.3299 | 14,145 | 34,071 | 23,388 |
| | | | | Indel | 0.0314 | 0.0372 | 0.3863 | 0.0679 | 226 | 5,844 | 359 |
| | | | 0.4 | SNV | 0.0475 | 0.1281 | 0.1688 | 0.1456 | 6,334 | 43,124 | 31,199 |
| | | | | Indel | 0.0169 | 0.0179 | 0.1983 | 0.0328 | 116 | 6,370 | 469 |
| | | | 0.2 | SNV | 0.0144 | 0.0557 | 0.0690 | 0.0617 | 2,591 | 43,893 | 34,942 |
| | | | | Indel | 0.0115 | 0.0047 | 0.0838 | 0.0089 | 49 | 10,438 | 536 |
| | DeepSomatic | 50x | 1.0 | SNV | 0.6334 | 0.5993 | 0.9157 | 0.7245 | 34,368 | 22,978 | 3,165 |
| | | | | Indel | 0.0992 | 0.1030 | 0.1350 | 0.1169 | 79 | 688 | 506 |
| | | | 0.8 | SNV | 0.5877 | 0.5805 | 0.9096 | 0.7087 | 34,139 | 24,671 | 3,394 |
| | | | | Indel | 0.1038 | 0.0993 | 0.2325 | 0.1392 | 136 | 1,233 | 449 |
| | | | 0.6 | SNV | 0.4980 | 0.5293 | 0.8929 | 0.6647 | 33,515 | 29,802 | 4,018 |
| | | | | Indel | 0.0801 | 0.0879 | 0.1385 | 0.1075 | 81 | 841 | 504 |
| | | | 0.4 | SNV | 0.3992 | 0.4823 | 0.8758 | 0.6221 | 32,870 | 35,278 | 4,663 |
| | | | | Indel | 0.0435 | 0.0429 | 0.1282 | 0.0643 | 75 | 1,674 | 510 |
| | | | 0.2 | SNV | 0.2878 | 0.4096 | 0.7586 | 0.5320 | 28,471 | 41,035 | 9,062 |
| | | | | Indel | 0.0240 | 0.0297 | 0.0308 | 0.0302 | 18 | 588 | 567 |

b

| Dataset | Caller | Tumor coverage | Tumor purity | w/ Verdict | Type | Precision | Recall | F1-score | TP | FP | FN |
|---|---|---|---|---|---|---|---|---|---|---|---|
| COLO829 | ClairS-TO SS | 50x | 0.6 | ✓ | SNV | 0.6642 | 0.8579 | 0.7488 | 32,200 | 16,277 | 5,333 |
| | | | | | Indel | 0.1990 | 0.3350 | 0.2497 | 196 | 789 | 389 |
| | | | | × | SNV | 0.6048 | 0.9204 | 0.7299 | 34,544 | 22,571 | 2,989 |
| | | | | | Indel | 0.1491 | 0.4667 | 0.2260 | 273 | 1,558 | 312 |
| | | | 0.4 | ✓ | SNV | 0.6845 | 0.8242 | 0.7479 | 30,936 | 14,261 | 6,597 |
| | | | | | Indel | 0.1349 | 0.4205 | 0.2042 | 246 | 1,578 | 339 |
| | | | | × | SNV | 0.5886 | 0.8767 | 0.7043 | 32,906 | 23,001 | 4,627 |
| | | | | | Indel | 0.0952 | 0.4376 | 0.1564 | 256 | 2,433 | 329 |
| | | | 0.2 | ✓ | SNV | 0.6318 | 0.7102 | 0.6687 | 26,655 | 15,532 | 10,878 |
| | | | | | Indel | 0.0596 | 0.3248 | 0.1007 | 190 | 2,999 | 395 |
| | | | | × | SNV | 0.4971 | 0.7426 | 0.5956 | 27,873 | 28,197 | 9,660 |
| | | | | | Indel | 0.0393 | 0.3350 | 0.0703 | 196 | 4,794 | 389 |
| | ClairS-TO SSRS | 50x | 0.6 | ✓ | SNV | 0.6695 | 0.8560 | 0.7514 | 32,128 | 15,857 | 5,405 |
| | | | | | Indel | 0.1942 | 0.3573 | 0.2517 | 209 | 867 | 376 |
| | | | | × | SNV | 0.6084 | 0.9170 | 0.7315 | 34,418 | 22,157 | 3,115 |
| | | | | | Indel | 0.1484 | 0.3812 | 0.2136 | 223 | 1,280 | 362 |
| | | | 0.4 | ✓ | SNV | 0.6862 | 0.8255 | 0.7495 | 30,984 | 14,166 | 6,549 |
| | | | | | Indel | 0.1552 | 0.2462 | 0.1904 | 144 | 784 | 441 |
| | | | | × | SNV | 0.5899 | 0.8781 | 0.7057 | 32,958 | 22,911 | 4,575 |
| | | | | | Indel | 0.1010 | 0.2632 | 0.1460 | 154 | 1,370 | 431 |
| | | | 0.2 | ✓ | SNV | 0.6992 | 0.6430 | 0.6699 | 24,132 | 10,384 | 13,401 |
| | | | | | Indel | 0.0890 | 0.0855 | 0.0872 | 50 | 512 | 535 |
| | | | | × | SNV | 0.5268 | 0.6752 | 0.5918 | 25,342 | 22,765 | 12,191 |
| | | | | | Indel | 0.0371 | 0.0906 | 0.0526 | 53 | 1,377 | 532 |

#### Supplementary Table 5. Performance on ONT COLO829 dataset across genomic contexts

The results of ClairS-TO (SS and SSRS model) and DeepSomatic for SNV and Indel on 50-fold coverage of ONT COLO829 across different challenging genomic contexts, within five categories: "Low complexity", "Segmental duplications", "Low mappability", "Functional regions", and "Other difficult regions", defined by GIAB Stratifications v3.3.

| Dataset | Caller | Tumor coverage | Stratification type | Stratification subtype | Type | Precision | Recall | F1-score | TP | FP | FN |
|---|---|---|---|---|---|---|---|---|---|---|---|
| COLO829 | ClairS-TO SS | 50x | Low complexity | Homopol_4-6bp | SNV | 0.6947 | 0.9321 | 0.7960 | 11,200 | 4,923 | 816 |
| | | | | | Indel | 0.2063 | 0.5495 | 0.3000 | 111 | 427 | 91 |
| | | | | Homopol_7-11bp | SNV | 0.6164 | 0.8503 | 0.7147 | 625 | 389 | 110 |
| | | | | | Indel | 0.1728 | 0.2240 | 0.1951 | 28 | 134 | 97 |
| | | | | Homopol_ge12bp | SNV | 0.3828 | 0.4969 | 0.4324 | 80 | 129 | 81 |
| | | | | | Indel | 0.0000 | 0.0000 | 0.0000 | 0 | 0 | 46 |
| | | | | Imp_Homopol_ge11bp | SNV | 0.5511 | 0.7537 | 0.6366 | 410 | 334 | 134 |
| | | | | | Indel | 0.1071 | 0.0349 | 0.0526 | 3 | 25 | 83 |
| | | | | TR_le50bp | SNV | 0.6093 | 0.8479 | 0.7091 | 262 | 168 | 47 |
| | | | | | Indel | 0.1739 | 0.1600 | 0.1667 | 4 | 19 | 21 |
| | | | | TR51-220bp | SNV | 0.6323 | 0.8598 | 0.7287 | 282 | 164 | 46 |
| | | | | | Indel | 0.1034 | 0.1200 | 0.1111 | 3 | 26 | 22 |
| | | | | TR201-10kbp | SNV | 0.5821 | 0.7130 | 0.6409 | 390 | 280 | 157 |
| | | | | | Indel | 0.0882 | 0.0909 | 0.0896 | 3 | 31 | 30 |
| | | | | TR_ge101bp | SNV | 0.6105 | 0.6377 | 0.6238 | 718 | 458 | 408 |
| | | | | | Indel | 0.0889 | 0.0851 | 0.0870 | 4 | 41 | 43 |
| | | | | TR_and_Homopol | SNV | 0.5696 | 0.7917 | 0.6625 | 2,170 | 1,640 | 571 |
| | | | | | Indel | 0.0918 | 0.1444 | 0.1122 | 39 | 386 | 231 |
| | | | Segmental duplications | SegDups | SNV | 0.5998 | 0.6635 | 0.6301 | 1,262 | 842 | 640 |
| | | | | | Indel | 0.1170 | 0.3548 | 0.1760 | 11 | 83 | 20 |
| | | | | SegDups_gt10kb | SNV | 0.5983 | 0.6346 | 0.6160 | 1,077 | 723 | 620 |
| | | | | | Indel | 0.1273 | 0.2593 | 0.1707 | 7 | 48 | 20 |
| | | | Low mappability | LowMap | SNV | 0.5903 | 0.6899 | 0.6362 | 1,686 | 1,170 | 758 |
| | | | | | Indel | 0.1383 | 0.3095 | 0.1912 | 13 | 81 | 29 |
| | | | Other difficult regions | L1H | SNV | 0.4943 | 0.9149 | 0.6418 | 43 | 44 | 4 |
| | | | | | Indel | 1.0000 | 1.0000 | 1.0000 | 1 | 0 | 0 |
| | | | | MHC | SNV | 0.7901 | 0.8649 | 0.8258 | 64 | 17 | 10 |
| | | | | | Indel | 0.0000 | 0.0000 | 0.0000 | 0 | 9 | 0 |
| | | | Functional regions | CDS | SNV | 0.7188 | 0.9186 | 0.8065 | 271 | 106 | 24 |
| | | | | | Indel | 0.4167 | 1.0000 | 0.5882 | 5 | 7 | 0 |
| COLO829 | ClairS-TO SSRS | 50x | Low complexity | Homopol_4-6bp | SNV | 0.7163 | 0.9204 | 0.8057 | 11,060 | 4,380 | 956 |
| | | | | | Indel | 0.2043 | 0.5644 | 0.3000 | 114 | 444 | 88 |
| | | | | Homopol_7-11bp | SNV | 0.6318 | 0.8476 | 0.7240 | 623 | 363 | 112 |
| | | | | | Indel | 0.2102 | 0.2960 | 0.2458 | 37 | 139 | 88 |
| | | | | Homopol_ge12bp | SNV | 0.3952 | 0.5155 | 0.4474 | 83 | 127 | 78 |
| | | | | | Indel | 0.0000 | 0.0000 | 0.0000 | 0 | 0 | 46 |
| | | | | Imp_Homopol_ge11bp | SNV | 0.5726 | 0.7463 | 0.6480 | 406 | 303 | 138 |
| | | | | | Indel | 0.2500 | 0.0698 | 0.1091 | 6 | 18 | 80 |
| | | | | TR_le50bp | SNV | 0.6072 | 0.8706 | 0.7154 | 269 | 174 | 40 |
| | | | | | Indel | 0.1600 | 0.1600 | 0.1600 | 4 | 21 | 21 |
| | | | | TR51-220bp | SNV | 0.6453 | 0.8598 | 0.7373 | 282 | 155 | 46 |
| | | | | | Indel | 0.1364 | 0.1200 | 0.1277 | 3 | 19 | 22 |
| | | | | TR201-10kbp | SNV | 0.5780 | 0.7788 | 0.6636 | 426 | 311 | 121 |
| | | | | | Indel | 0.0556 | 0.0606 | 0.0580 | 2 | 34 | 31 |
| | | | | TR_ge101bp | SNV | 0.5985 | 0.7043 | 0.6471 | 793 | 532 | 333 |
| | | | | | Indel | 0.0556 | 0.0851 | 0.0672 | 4 | 68 | 43 |
| | | | | TR_and_Homopol | SNV | 0.6092 | 0.7775 | 0.6831 | 2,131 | 1,367 | 610 |
| | | | | | Indel | 0.1494 | 0.1704 | 0.1592 | 46 | 262 | 224 |
| | | | Segmental duplications | SegDups | SNV | 0.5918 | 0.6951 | 0.6393 | 1,322 | 912 | 580 |
| | | | | | Indel | 0.1951 | 0.2581 | 0.2222 | 8 | 33 | 23 |
| | | | | SegDups_gt10kb | SNV | 0.5796 | 0.6735 | 0.6231 | 1,143 | 829 | 554 |
| | | | | | Indel | 0.1951 | 0.2963 | 0.2353 | 8 | 33 | 19 |
| | | | Low mappability | LowMap | SNV | 0.5889 | 0.7197 | 0.6478 | 1,759 | 1,228 | 685 |
| | | | | | Indel | 0.1340 | 0.3095 | 0.1871 | 13 | 84 | 29 |
| | | | Other difficult regions | L1H | SNV | 0.6444 | 0.6170 | 0.6304 | 29 | 16 | 18 |
| | | | | | Indel | 0.3333 | 1.0000 | 0.5000 | 1 | 2 | 0 |
| | | | | MHC | SNV | 0.7901 | 0.8649 | 0.8258 | 64 | 17 | 10 |
| | | | | | Indel | 0.0000 | 0.0000 | 0.0000 | 0 | 1 | 0 |
| | | | Functional regions | CDS | SNV | 0.7535 | 0.9017 | 0.8210 | 266 | 87 | 29 |
| | | | | | Indel | 0.3846 | 1.0000 | 0.5556 | 5 | 8 | 0 |
| COLO829 | DeepSomatic | 50x | Low complexity | Homopol_4-6bp | SNV | 0.6542 | 0.9127 | 0.7621 | 10,967 | 5,797 | 1,049 |
| | | | | | Indel | 0.0776 | 0.2327 | 0.1163 | 47 | 559 | 155 |
| | | | | Homopol_7-11bp | SNV | 0.4819 | 0.8490 | 0.6148 | 624 | 671 | 111 |
| | | | | | Indel | 0.0598 | 0.1200 | 0.0798 | 15 | 236 | 110 |
| | | | | Homopol_ge12bp | SNV | 0.2096 | 0.4596 | 0.2879 | 74 | 279 | 87 |
| | | | | | Indel | 0.0000 | 0.0000 | 0.0000 | 0 | 174 | 46 |
| | | | | Imp_Homopol_ge11bp | SNV | 0.4296 | 0.6452 | 0.5158 | 351 | 466 | 193 |
| | | | | | Indel | 0.0256 | 0.0116 | 0.0160 | 1 | 38 | 85 |
| | | | | TR_le50bp | SNV | 0.5207 | 0.8544 | 0.6471 | 264 | 243 | 45 |
| | | | | | Indel | 0.0638 | 0.1200 | 0.0833 | 3 | 44 | 22 |
| | | | | TR51-220bp | SNV | 0.4699 | 0.7622 | 0.5814 | 250 | 282 | 78 |
| | | | | | Indel | 0.0222 | 0.0400 | 0.0286 | 1 | 44 | 24 |
| | | | | TR201-10kbp | SNV | 0.4209 | 0.6033 | 0.4959 | 330 | 454 | 217 |
| | | | | | Indel | 0.0769 | 0.0303 | 0.0435 | 1 | 12 | 32 |
| | | | | TR_ge101bp | SNV | 0.4102 | 0.6128 | 0.4915 | 690 | 992 | 436 |
| | | | | | Indel | 0.0465 | 0.0426 | 0.0444 | 2 | 41 | 45 |
| | | | | TR_and_Homopol | SNV | 0.4417 | 0.7209 | 0.5477 | 1,976 | 2,498 | 765 |
| | | | | | Indel | 0.0395 | 0.0444 | 0.0418 | 12 | 292 | 258 |
| | | | Segmental duplications | SegDups | SNV | 0.3977 | 0.6467 | 0.4925 | 1,230 | 1,863 | 672 |
| | | | | | Indel | 0.0197 | 0.1935 | 0.0357 | 6 | 299 | 25 |
| | | | | SegDups_gt10kb | SNV | 0.3792 | 0.6299 | 0.4734 | 1,069 | 1,750 | 628 |
| | | | | | Indel | 0.0213 | 0.0741 | 0.0331 | 2 | 92 | 25 |
| | | | Low mappability | LowMap | SNV | 0.3881 | 0.6825 | 0.4948 | 1,668 | 2,630 | 776 |
| | | | | | Indel | 0.0526 | 0.0952 | 0.0678 | 4 | 72 | 38 |
| | | | Other difficult regions | L1H | SNV | 0.3009 | 0.7234 | 0.4250 | 34 | 79 | 13 |
| | | | | | Indel | 1.0000 | 1.0000 | 1.0000 | 1 | 0 | 0 |
| | | | | MHC | SNV | 0.6292 | 0.7568 | 0.6871 | 56 | 33 | 18 |
| | | | | | Indel | 0.0000 | 0.0000 | 0.0000 | 0 | 4 | 0 |
| | | | Functional regions | CDS | SNV | 0.6522 | 0.9153 | 0.7616 | 270 | 144 | 25 |
| | | | | | Indel | 0.1429 | 0.4000 | 0.2105 | 2 | 12 | 3 |

#### Supplementary Table 6. The reasons of 300 FPs and 300 FNs

The table is in the file named "Supplementary_table_6.xlsx". The FPs and FNs were randomly picked from the variant called using ClairS-TO SS model on 50-fold coverage of ONT COLO829.

#### Supplementary Table 7. Performance on ONT COLO829 dataset with different ablation settings

(a) The performance of SNV and Indel on 50-fold coverage of ONT COLO829 using 1) ClairS-TO SS, 2) RS, and 3) SSRS models. (b) The performance of SNV and Indel on 50-fold coverage of ONT COLO829 using 1) both the AFF & NEG networks, 2) only the AFF network, and 3) only the NEG network, from the ClairS-TO SSRS model. (c) The performance of using ClairS-TO SS or ClairS-TO SSRS model with different PoNs or combinations for filtering.

**a**

| Dataset | Caller | Tumor coverage | Type | AUPRC | Precision | Recall | F1-score | TP | FP | FN |
|---|---|---|---|---|---|---|---|---|---|---|
| COLO829 | ClairS-TO SS | 50x | SNV | 0.6176 | 0.6486 | 0.9282 | 0.7636 | 34,837 | 18,870 | 2,696 |
| | | | Indel | 0.1833 | 0.1963 | 0.4889 | 0.2801 | 286 | 1,171 | 299 |
| | ClairS-TO RS | 50x | SNV | 0.6582 | 0.6686 | 0.8986 | 0.7667 | 33,728 | 16,720 | 3,805 |
| | | | Indel | 0.1889 | 0.1874 | 0.3521 | 0.2447 | 206 | 893 | 379 |
| | ClairS-TO SSRS | 50x | SNV | 0.6634 | 0.6614 | 0.9163 | 0.7683 | 34,391 | 17,605 | 3,142 |
| | | | Indel | 0.2019 | 0.1974 | 0.4957 | 0.2824 | 290 | 1,179 | 295 |

**b**

| Dataset | Caller | Neural network | Tumor coverage | Type | AUPRC | Precision | Recall | F1-score | TP | FP | FN |
|---|---|---|---|---|---|---|---|---|---|---|---|
| COLO829 | ClairS-TO SSRS | AFF & NEG | 50x | SNV | 0.6634 | 0.6614 | 0.9163 | 0.7683 | 34,391 | 17,605 | 3,142 |
| | | | | Indel | 0.2019 | 0.1974 | 0.4957 | 0.2824 | 290 | 1,179 | 295 |
| | | AFF only | | SNV | 0.6363 | 0.6489 | 0.9284 | 0.7639 | 34,847 | 18,854 | 2,686 |
| | | | | Indel | 0.1999 | 0.1933 | 0.5094 | 0.2802 | 298 | 1,244 | 287 |
| | | NEG only | | SNV | 0.6562 | 0.6508 | 0.9265 | 0.7646 | 34,774 | 18,656 | 2,759 |
| | | | | Indel | 0.2005 | 0.1859 | 0.4786 | 0.2678 | 280 | 1,226 | 305 |

**c**

| Dataset | Caller | Tumor coverage | PoNs | Type | Precision | Recall | F1-score | TP | FP | FN |
|---|---|---|---|---|---|---|---|---|---|---|
| COLO829 | ClairS-TO SS | 50x | None | SNV | 0.0106 | 0.9111 | 0.0211 | 34,195 | 3,176,607 | 3,338 |
| | | | | Indel | 0.0008 | 0.7282 | 0.0017 | 426 | 512,212 | 159 |
| | | | gnomAD | SNV | 0.2252 | 0.8759 | 0.3583 | 32,875 | 113,104 | 4,658 |
| | | | | Indel | 0.0023 | 0.5675 | 0.0047 | 332 | 141,197 | 253 |
| | | | dbSNP | SNV | 0.2669 | 0.8692 | 0.4084 | 32,624 | 89,590 | 4,909 |
| | | | | Indel | 0.0041 | 0.5214 | 0.0081 | 305 | 74,748 | 280 |
| | | | 1000g PoN | SNV | 0.0180 | 0.9110 | 0.0354 | 34,194 | 1,862,433 | 3,339 |
| | | | | Indel | 0.0011 | 0.6427 | 0.0023 | 376 | 329,544 | 209 |
| | | | CoLoRSdb | SNV | 0.5713 | 0.8937 | 0.6970 | 33,545 | 25,171 | 3,988 |
| | | | | Indel | 0.0754 | 0.3573 | 0.1246 | 209 | 2,562 | 376 |
| | | | All | SNV | 0.6601 | 0.8842 | 0.7559 | 33,186 | 17,090 | 4,347 |
| | | | | Indel | 0.1029 | 0.3538 | 0.1595 | 207 | 1,804 | 378 |
| | ClairS-TO SSRS | 50x | None | SNV | 0.0118 | 0.6383 | 0.0231 | 23,958 | 2,014,183 | 13,575 |
| | | | | Indel | 0.0023 | 0.0479 | 0.0043 | 28 | 12,340 | 557 |
| | | | gnomAD | SNV | 0.2574 | 0.8550 | 0.3956 | 32,090 | 92,600 | 5,443 |
| | | | | Indel | 0.0121 | 0.0906 | 0.0213 | 53 | 4,332 | 532 |
| | | | dbSNP | SNV | 0.2996 | 0.8874 | 0.4480 | 33,308 | 77,855 | 4,225 |
| | | | | Indel | 0.0250 | 0.0906 | 0.0392 | 53 | 2,067 | 532 |
| | | | 1000g PoN | SNV | 0.0203 | 0.4170 | 0.0387 | 15,651 | 756,069 | 21,882 |
| | | | | Indel | 0.0030 | 0.0479 | 0.0056 | 28 | 9,349 | 557 |
| | | | CoLoRSdb | SNV | 0.5760 | 0.9316 | 0.7119 | 34,966 | 25,737 | 2,567 |
| | | | | Indel | 0.1131 | 0.4718 | 0.1825 | 276 | 2,164 | 309 |
| | | | All | SNV | 0.6404 | 0.9495 | 0.7649 | 35639 | 20012 | 1894 |
| | | | | Indel | 0.1514 | 0.4701 | 0.2291 | 275 | 1,541 | 310 |

#### Supplementary Table 8. Performance on PacBio and Illumina data

(a) The performance of ClairS-TO (SS and SSRS model), Clair3, and DeepSomatic on 50-fold coverage of PacBio Revio COLO829. (b) The performance of ClairS-TO (SS and SSRS model), DeepSomatic, Mutect2, Octopus, and Pisces on 50-fold coverage of Illumina COLO829.

**a**

| Dataset | Caller | Tumor coverage | Type | AUPRC | Precision | Recall | F1-score | TP | FP | FN |
|---|---|---|---|---|---|---|---|---|---|---|
| COLO829 | ClairS-TO SS | 50x | SNV | 0.6439 | 0.6814 | 0.9258 | 0.7850 | 36,185 | 16,917 | 2,901 |
| | | | Indel | 0.1886 | 0.1779 | 0.4951 | 0.2617 | 302 | 1,396 | 308 |
| | ClairS-TO SSRS | 50x | SNV | 0.6667 | 0.6840 | 0.9248 | 0.7864 | 36,148 | 16,697 | 2,938 |
| | | | Indel | 0.1972 | 0.1864 | 0.4410 | 0.2621 | 269 | 1,174 | 341 |
| | Clair3 | 50x | SNV | 0.5576 | 0.5305 | 0.8502 | 0.6533 | 33,231 | 29,409 | 5,855 |
| | | | Indel | 0.1365 | 0.1359 | 0.3820 | 0.2004 | 233 | 1,482 | 377 |
| | DeepSomatic | 50x | SNV | 0.6588 | 0.6382 | 0.8991 | 0.7465 | 35,143 | 19,920 | 3,943 |
| | | | Indel | 0.1420 | 0.1334 | 0.2016 | 0.1606 | 123 | 799 | 487 |

**b**

| Dataset | Caller | Tumor coverage | Type | AUPRC | Precision | Recall | F1-score | TP | FP | FN |
|---|---|---|---|---|---|---|---|---|---|---|
| COLO829 | ClairS-TO SS | 50x | SNV | 0.6228 | 0.6721 | 0.8877 | 0.7650 | 35,712 | 17,421 | 4,520 |
| | | | Indel | 0.2422 | 0.2148 | 0.4629 | 0.2934 | 331 | 1,210 | 384 |
| | ClairS-TO SSRS | 50x | SNV | 0.6500 | 0.6873 | 0.8751 | 0.7699 | 35,207 | 16,017 | 5,025 |
| | | | Indel | 0.2334 | 0.2151 | 0.4657 | 0.2943 | 333 | 1,215 | 382 |
| | DeepSomatic | 50x | SNV | 0.5039 | 0.6100 | 0.6280 | 0.6189 | 25,264 | 16,151 | 14,968 |
| | | | Indel | 0.0521 | 0.0533 | 0.1510 | 0.0788 | 108 | 1,919 | 607 |
| | Mutect2 | 50x | SNV | 0.3570 | 0.4603 | 0.7896 | 0.5816 | 31,768 | 37,243 | 8,464 |
| | | | Indel | 0.0223 | 0.0260 | 0.2801 | 0.0476 | 200 | 7,492 | 514 |
| | Octopus | 50x | SNV | 0.2119 | 0.2746 | 0.7566 | 0.4030 | 30,440 | 80,410 | 9,792 |
| | | | Indel | 0.0044 | 0.0050 | 0.1459 | 0.0096 | 104 | 20,809 | 609 |
| | Pisces | 50x | SNV | 0.0635 | 0.1290 | 0.8487 | 0.2240 | 34,145 | 230,502 | 6,085 |
| | | | Indel | 0.0008 | 0.0015 | 0.5930 | 0.0030 | 424 | 282,328 | 291 |

### Supplementary Methods

#### Pileup input

The pileup input of ClairS-TO comprises 1,122 integers (i.e., 33 positions with 34 features at each position). The 34 features provide information on the counts on both the forward and reverse strand of 1) nucleotides, 2) insertions, 3) deletions, 4) nucleotides with low mapping quality, and 5) nucleotides with low base quality. Below is a detailed breakdown of each feature:

1-4: A+/C+/G+/T+: The counts of A/C/G/T nucleotides present in the forward strand.

5: $I_S$+: The count of insertions that have the same starting positions as the candidate site in the forward strand.

6: $I_{1S}$+: Similar to $I_S$+ but only counting the insertion with the highest read support.

7: $D_S$+: The count of deletions that have the same starting positions as the candidate site in the forward strand.

8: $D_{1S}$+: Similar to $D_S$+ but only counting the deletion with the highest read support.

9: $D_R$+: The count of all deletions in the forward strand that crossed the position except for the first base of each deletion.

10-13: A-/C-/G-/T-: The counts of A/C/G/T nucleotides present in the reverse strand.

14: $I_S$-: The count of insertions that have the same starting positions as the candidate site in the reverse strand.

15: $I_{1S}$-: Similar to $I_S$- but only counting the insertion with the highest read support.

16: $D_S$-: The count of deletions that have the same starting positions as the candidate site in the reverse strand.

17: $D_{1S}$-: Similar to $D_S$- but only counting the deletion with the highest read support.

18: $D_R$-: The count of all deletions in the reverse strand that crossed the position except for the first base of each deletion.

19-22: $A_{LMQ}$+/$C_{LMQ}$+/$G_{LMQ}$+/$T_{LMQ}$+: The counts of A/C/G/T nucleotides with low mapping quality (MQ<20) in the forward strand.

23-26: $A_{LBQ}$+/$C_{LBQ}$+/$G_{LBQ}$+/$T_{LBQ}$+: The counts of A/C/G/T nucleotides with low base quality (BQ<30) in the forward strand.

27-30: $A_{LMQ}$-/$C_{LMQ}$-/$G_{LMQ}$-/$T_{LMQ}$-: The counts of A/C/G/T nucleotides with low mapping quality (MQ<20) in the reverse strand.

31-34: $A_{LBQ}$-/$C_{LBQ}$-/$G_{LBQ}$-/$T_{LBQ}$-: The counts of A/C/G/T nucleotides with low base quality (BQ<30) in the reverse strand.

### Command lines used

#### Read alignment

##### Minimap2 (v2.17-r941)

```
# Align ONT reads using minimap2 to GRCh38_no_alt by default
minimap2 -t ${THREADS} -aL -z 600,200 -x map-ont ref.fa input.fastq.gz | samtools view -bh
-o output.unsorted.bam -
samtools sort -@${THREADS} -o output.sorted.bam output.unsorted.bam && samtools index -@
${THREADS} output.sorted.bam
```

#### BAM subsampling

##### Samtools (v1.15.1)

```
samtools view -@ ${THREADS} -s ${RATIO}.${RATIO} -b -o subsampled.bam ${BAM}
samtools index -@ ${THREADS} subsampled.bam
```

#### Coverage calculation

##### Mosdepth (v0.3.1)

```
mosdepth -t ${THREADS} -n -x --quantize 0:15:150: output ${BAM}
```

#### Alignment statistical summary

##### NanoPlot (v1.40.2)

```
NanoPlot -t ${THREADS} --bam ${BAM} --N50 -o bamplots
```

#### Generating BAMs with different tumor purities

```
# Use ${TUMOR_PURITY} to add tumor purity
pypy3 clairs_to.py gen_contaminated_bam \
       --tumor_bam_fn ${TUMOR_BAM_FILE_PATH} \
       --normal_bam_fn ${NORMAL_BAM_FILE_PATH} \
       --tumor_purity ${TUMOR_PURITY} \
       --output_dir ${OUTPUT_DIR} \
       --tumor_bam_coverage ${TUMOR_BAM_COVERAGE} \
       --normal_bam_coverage ${NORMAL_BAM_COVERAGE} \
       --mosdepth ${MOSDEPTH_PATH}
```

#### Running ClairS-TO (v0.4.0)

```
run_clairs_to \
       --tumor_bam_fn ${TUMOR_BAM_FILE_PATH} \
       --ref_fn ${REF} \
       --threads ${THREADS} \
       --platform ${PLATFORM} \
       --output_dir ${OUTPUT_DIR}
```

#### Running Clair3 (v1.0.8)

```
# The Clair3 R10.4.1 Q20+ model was downloaded from https://github.com/nanoporetech/rerio
# maintained by ONT developer
# Then run the following command in tumor BAM
docker run -it \
  -v ${INPUT_DIR}:${INPUT_DIR} \
  -v ${OUTPUT_DIR}:${OUTPUT_DIR} \
  hkubal/clair3:v1.0.8 \
  /opt/bin/run_clair3.sh \
       --bam_fn=${INPUT_DIR}/input.bam \
       --ref_fn=${INPUT_DIR}/ref.fa \
       --threads=${THREADS} \
       --platform="${PLATFORM}" \
       --model_path="${INPUT_DIR}/r1041_e82_400bps_sup_v420" \
       --output=${OUTPUT_DIR}
```

#### Running other somatic variant callers for ONT data

##### DeepSomatic (v1.7.0)

```
docker run -it \
```

```
  -v ${INPUT_DIR}:${INPUT_DIR} \
  -v ${OUTPUT_DIR}:${OUTPUT_DIR} \
  google/deepsomatic:latest \
  run_deepsomatic \
       --model_type="ONT_TUMOR_ONLY" \
       --ref=${REF} \
       --reads_tumor=${TUMOR_BAM_FILE_PATH} \
       --output_vcf=${OUTPUT_DIR}/output.vcf.gz \
       --sample_name_tumor="tumor" \
       --num_shards=${THREADS} \
       --use_default_pon_filtering True
```

#### Running other somatic variant callers for PacBio data

##### DeepSomatic (v1.7.0)

```
docker run -it \
  -v ${INPUT_DIR}:${INPUT_DIR} \
  -v ${OUTPUT_DIR}:${OUTPUT_DIR} \
  google/deepsomatic:latest \
  run_deepsomatic \
       --model_type="PACBIO_TUMOR_ONLY" \
       --ref=${REF} \
       --reads_tumor=${TUMOR_BAM_FILE_PATH} \
       --output_vcf=${OUTPUT_DIR}/output.vcf.gz \
       --sample_name_tumor="tumor" \
       --num_shards=${THREADS} \
       --use_default_pon_filtering True
```

#### Running other somatic variant callers for Illumina data

##### DeepSomatic (v1.7.0)

```
docker run -it \
  -v ${INPUT_DIR}:${INPUT_DIR} \
  -v ${OUTPUT_DIR}:${OUTPUT_DIR} \
  google/deepsomatic:latest \
  run_deepsomatic \
       --model_type="WGS_TUMOR_ONLY" \
       --ref=${REF} \
       --reads_tumor=${TUMOR_BAM_FILE_PATH} \
       --output_vcf=${OUTPUT_DIR}/output.vcf.gz \
```

```
        --sample_name_tumor="tumor" \
        --num_shards=${THREADS} \
        --use_default_pon_filtering True
```

##### Mutect2 (v4.2.6.1)

```
gatk --java-options -Xmx12G Mutect2 \
        --reference ${REF} \
        --intervals ${BED_FILE_PATH} \
        --input ${TUMOR_BAM_FILE_PATH} \
        --output ${OUTPUT_DIR}/unfiltered.mutect2.vcf \
        --pon 1000g_pon.hg38.vcf.gz \
        --germline-resource af-only-gnomad.hg38.vcf.gz

gatk --java-options -Xmx12G FilterMutectCalls \
        --variant ${OUTPUT_DIR}/unfiltered.mutect2.vcf \
        --reference ${REF} \
        --output ${OUTPUT_DIR}/mutect2.vcf)
```

##### Octopus (v0.7.4)

```
octopus \
        -R ${REF} \
        -I ${TUMOR_BAM_FILE_PATH} \
        -C cancer \
        --threads ${THREADS} \
        -o ${OUTPUT_FOLDER}/octopus.vcf
```

##### Pisces (v5.3.0)

```
Pisces \
        -g ${REF} \
        -b ${TUMOR_BAM_FILE_PATH} \
        -MaxNumThreads ${THREADS} \
        -OutFolder ${OUTPUT_FOLDER}
```

#### Benchmarking

##### som.py (v0.3.12)

```
som.py ${SEQC2_BASELINE_VCF} output.vcf.gz \
        -T ${SEQC2_CONFIDENT_BED} \
        -f ${SEQC2_CONFIDENT_BED} \
        -r ${REF} \
        -o benchmark_result
```

Calculate Precision, Recall, F1-Score with "compare_vcf" submodule in ClairS-TO

```
pypy3 clairs_to.py compare_vcf \
       --truth_vcf_fn ${SEQC2_BASELINE_VCF} \
       --input_vcf_fn output.vcf.gz \
       --bed_fn ${SEQC2_CONFIDENT_BED} \
       --output_dir benchmark_result \
       --input_filter_tag 'PASS' \
       --min_qual ${MIN_QUAL} \
       --min_af ${MIN_AF} \
       --output_best_f1_score \
       --tumor_bam_fn ${TUMOR_BAM_FILE_PATH}
```

### Data availability

#### GIAB truth variants

HG001 (NA12878), GRCh38, v4.2.1

https://ftp-trace.ncbi.nlm.nih.gov/giab/ftp/release/NA12878_HG001/NISTv4.2.1/GRCh38/

HG002 (NA24385), GRCh38, v4.2.1

https://ftp-trace.ncbi.nlm.nih.gov/giab/ftp/release/AshkenazimTrio/HG002_NA24385_son/NISTv4.2.1/GRCh38/

HG003 (NA24149), GRCh38, v4.2.1

https://ftp-trace.ncbi.nlm.nih.gov/giab/ftp/release/AshkenazimTrio/HG003_NA24149_father/NISTv4.2.1/GRCh38/

HG004 (NA24143), GRCh38, v4.2.1

https://ftp-trace.ncbi.nlm.nih.gov/giab/ftp/release/AshkenazimTrio/HG004_NA24143_mother/NISTv4.2.1/GRCh38/

#### NYGC truth variants for COLO829 sample

COLO829, GRCh38

https://bioinformatics.nygenome.org/3-cancer-cell-lines-on-2-sequencers/

#### SEQC truth variants for HCC1395 sample

HCC1395, GRCh38, v1.2

https://ftp-trace.ncbi.nlm.nih.gov/ReferenceSamples/seqc/Somatic_Mutation_WG/release/v1.2

#### Four real cancer cell-lines variants from Park et al.

HCC1937, GRCh38

https://storage.googleapis.com/brain-genomics-public/publications/park2024_deepsomatic/benchmarking/DeepSomatic_multicancer-model_benchmark/vcfs/somatic_only/HCC1937_DeepSomatic_multicancer-model_somatic-only.vcf.gz

HCC1954, GRCh38

https://storage.googleapis.com/brain-genomics-public/publications/park2024_deepsomatic/benchmarking/DeepSomatic_multicancer-model_benchmark/vcfs/somatic_only/HCC1954_DeepSomatic_multicancer-model_somatic-only.vcf.gz

H1437, GRCh38

https://storage.googleapis.com/brain-genomics-public/publications/park2024_deepsomatic/benchmarking/DeepSomatic_multicancer-model_benchmark/vcfs/somatic_only/H1437_DeepSomatic_multicancer-model_somatic-only.vcf.gz

H2009, GRCh38

https://storage.googleapis.com/brain-genomics-public/publications/park2024_deepsomatic/benchmarking/DeepSomatic_multicancer-model_benchmark/vcfs/somatic_only/H2009_DeepSomatic_multicancer-model_somatic-only.vcf.gz

#### Reference genomes

GRCh38_no_alt

https://ftp.ncbi.nlm.nih.gov/genomes/all/GCA/000/001/405/GCA_000001405.15_GRCh38/seqs_for_alignment_pipelines.ucsc_ids/GCA_000001405.15_GRCh38_no_alt_analysis_set.fna.gz

GRCh38 d1.vd1

https://ftp-trace.ncbi.nlm.nih.gov/ReferenceSamples/seqc/Somatic_Mutation_WG/technical/reference_genome/GRCh38/GRCh38.d1.vd1.fa

GRCh38 Stratification regions (v3.3)

https://ftp-trace.ncbi.nlm.nih.gov/giab/ftp/release/genome-stratifications/v3.3/GRCh38@all

#### ONT Sequencing Data

ONT EPI2ME Labs HG001 R10.4.1 Q20+, GRCh38_no_alt, 78.89-fold

https://labs.epi2me.io/giab-2023.05/

ONT EPI2ME Labs HG002 R10.4.1 Q20+, GRCh38_no_alt, 91.18-fold

https://labs.epi2me.io/giab-2023.05/

Park et al. HCC1937 R10.4.1 Q20+, GRCh38_no_alt, 158.14-fold https://storage.googleapis.com/brain-genomics-public/publications/park2024_deepsomatic/bams/ONT/minimap2_grch38_bams/minimap2_grch38_bams/1937_Tumor_ONT.GRCh38.sorted.bam

Park et al. HCC1954 R10.4.1 Q20+, GRCh38_no_alt, 82.44-fold

https://storage.googleapis.com/brain-genomics-public/publications/park2024_deepsomatic/bams/ONT/minimap2_grch38_bams/minimap2_grch38_bams/1954_Tumor_ONT.GRCh38.sorted.bam

Park et al. H1437 R10.4.1 Q20+, GRCh38_no_alt, 79.10-fold

https://storage.googleapis.com/brain-genomics-public/publications/park2024_deepsomatic/bams/ONT/minimap2_grch38_bams/minimap2_grch38_bams/1437_Tumor_ONT.GRCh38.sorted.bam

Park et al. H2009 R10.4.1 Q20+, GRCh38_no_alt, 106.92-fold

https://storage.googleapis.com/brain-genomics-public/publications/park2024_deepsomatic/bams/ONT/minimap2_grch38_bams/minimap2_grch38_bams/2009_Tumor_ONT.GRCh38.sorted.bam

Park et al. HCC1395 R10.4.1 Q20+, GRCh38_no_alt, 79.52-fold

https://storage.googleapis.com/brain-genomics-public/publications/park2024_deepsomatic/bams/ONT/minimap2_grch38_bams/minimap2_grch38_bams/1395_Tumor_ONT.GRCh38.sorted.bam

ONT EPI2ME Labs COLO829 R10.4.1 Q20+, GRCh38_no_alt, 101.80-fold

https://labs.epi2me.io/colo-2024.03/

#### PacBio Sequencing Data

PacBio HG003 HiFi Revio, GRCh38_no_alt, 61.15-fold

https://downloads.pacbcloud.com/public/revio/2022Q4/HG003-rep1/

PacBio HG004 HiFi Revio, GRCh38_no_alt, 62.67-fold

https://downloads.pacbcloud.com/public/revio/2022Q4/HG004-rep1/

Park et al. HCC1937 HiFi Revio, GRCh38_no_alt, 59.50-fold

https://storage.googleapis.com/brain-genomics-public/publications/park2024_deepsomatic/bams/HiFi/pbmm2_grch38_bams/1937_Tumor_HiFi.GRCh38.sorted.bam

Park et al. HCC1954 HiFi Revio, GRCh38_no_alt, 63.07-fold

https://storage.googleapis.com/brain-genomics-public/publications/park2024_deepsomatic/bams/HiFi/pbmm2_grch38_bams/1954_Tumor_HiFi.GRCh38.sorted.bam

Park et al. H1437 HiFi Revio, GRCh38_no_alt, 64.06-fold

https://storage.googleapis.com/brain-genomics-public/publications/park2024_deepsomatic/bams/HiFi/pbmm2_grch38_bams/1437_Tumor_HiFi.GRCh38.sorted.bam

Park et al. H2009 HiFi Revio, GRCh38_no_alt, 64.89-fold

https://storage.googleapis.com/brain-genomics-public/publications/park2024_deepsomatic/bams/HiFi/pbmm2_grch38_bams/2009_Tumor_HiFi.GRCh38.sorted.bam

PacBio COLO829 HiFi Revio, GRCh38_no_alt, 60.70-fold

https://downloads.pacbcloud.com/public/revio/2023Q2/COLO829/COLO829/

#### Illumina Sequencing Data

HG003 NovaSeq 6000 (NA24149), GRCh38_no_alt, 47.38-fold

https://storage.googleapis.com/brain-genomics-public/research/sequencing/grch38/bam/novaseq/wgs_pcr_free/50x/HG003.novaseq.pcr-free.50x.dedup.grch38.bam

HG004 NovaSeq 6000 (NA24385), GRCh38_no_alt, 46.36-fold

https://storage.googleapis.com/brain-genomics-public/research/sequencing/grch38/bam/novaseq/wgs_pcr_free/50x/HG004.novaseq.pcr-free.50x.dedup.grch38.bam

HG003 HiSeqX (NA24149), GRCh38_no_alt 43.77-fold

https://storage.googleapis.com/brain-genomics-public/research/sequencing/grch38/bam/hiseqx/wgs_pcr_free/40x/HG003.hiseqx.pcr-free.40x.dedup.grch38.bam

HG004 HiSeqX (NA24385), GRCh38_no_alt, 42.13-fold

https://storage.googleapis.com/brain-genomics-public/research/sequencing/grch38/bam/hiseqx/wgs_pcr_free/40x/HG004.hiseqx.pcr-free.40x.dedup.grch38.bam

Park et al. HCC1937 NovaSeq 6000, GRCh38, 256.28-fold

https://storage.googleapis.com/brain-genomics-public/publications/park2024_deepsomatic/bams/illumina/bwa_mem2_grch38_bams/bwa_mem2_grch38_bams/1937_Tumor_Illumina.GRCh38.sorted.bam

Park et al. HCC1954 NovaSeq 6000, GRCh38, 139.02-fold

https://storage.googleapis.com/brain-genomics-public/publications/park2024_deepsomatic/bams/illumina/bwa_mem2_grch38_bams/bwa_mem2_grch38_bams/1954_Tumor_Illumina.GRCh38.sorted.bam

Park et al. H1437 NovaSeq 6000, GRCh38, 203.94-fold

https://storage.googleapis.com/brain-genomics-public/publications/park2024_deepsomatic/bams/illumina/bwa_mem2_grch38_bams/bwa_mem2_grch38_bams/1437_Tumor_Illumina.GRCh38.sorted.bam

Park et al. H2009 NovaSeq 6000, GRCh38, 230.57-fold

https://storage.googleapis.com/brain-genomics-public/publications/park2024_deepsomatic/bams/illumina/bwa_mem2_grch38_bams/bwa_mem2_grch38_bams/2009_Tumor_Illumina.GRCh38.sorted.bam

COLO829 NovaSeq 6000, GRCh38, 234.44-fold

https://www.ncbi.nlm.nih.gov/projects/gap/cgi-bin/collection.cgi?study_id=phs000688.v1.p1
